# Linked mutations at adjacent nucleotides have shaped human population differentiation and protein evolution

**DOI:** 10.1101/329292

**Authors:** James G. D. Prendergast, Carys Pugh, Sarah E. Harris, David A. Hume, Ian J. Deary, Allan Beveridge

**Affiliations:** The Roslin Institute, University of Edinburgh, Easter Bush Campus, Midlothian, EH25 9RG, United Kingdom; Centre for Cognitive Ageing and Cognitive Epidemiology, Department of Psychology, The University of Edinburgh, Edinburgh, United Kingdom; The University of Edinburgh Centre for Genomic and Experimental Medicine and MRC Institute of Genetics and Molecular Medicine, Edinburgh, United Kingdom; Mater Research Institute-University of Queensland, Woolloongabba, Qld 4160, Australia; Glasgow Polyomics, College of Medical, Veterinary and Life Science, University of Glasgow, Glasgow, United Kingdom

**Keywords:** sequential dinucleotide mutations, multi-nucleotide polymorphisms, multi-nucleotide mutations, human mutation, DNA repair, epistatic selection, SDM, MNP

## Abstract

Despite the fundamental importance of single nucleotide polymorphisms (SNPs) to human evolution there are still large gaps in our understanding of the forces that shape their distribution across the genome. SNPs have been shown to not be distributed evenly, with directly adjacent SNPs found unusually frequently. Why this is the case is unclear. We illustrate how neighbouring SNPs that can’t be explained by a single mutation event (that we term here sequential dinucleotide mutations, SDMs) are driven by distinct mutational processes and selective pressures to SNPs and multinucleotide polymorphisms (MNPs). By studying variation across multiple populations, including a novel cohort of 1,358 Scottish genomes, we show that, SDMs are over twice as common as MNPs and like SNPs, display distinct mutational spectra across populations. These biases are though not only different to those observed among SNPs and MNPs, but also more divergent between human population groups. We show that the changes that make up SDMs are not independent, and identify a distinct mutational profile, CA → CG → TG, that is observed an order of magnitude more often than other SDMs, including others that involve the gain and subsequent deamination of CpG sites. This suggests these specific changes are driven by a distinct process. In coding regions particular SDMs are favoured, and especially those that lead to the creation of single codon amino acids. Intriguingly selection has favoured particular pathways through the amino acid code, with epistatic selection appearing to have disfavoured sequential non-synonymous changes.

## Introduction

Single nucleotide polymorphisms (SNPs) are the pillars of modern genetics studies (Gray et al., 2000; Brookes, 1999). From their use in genome-wide association studies to map the genetics of diseases (Bush and Moore, 2012), to studying patterns of evolution (Morin et al., 2004), SNPs are widely used and studied across fields. A common implicit assumption across these studies is that SNPs are independent (Schrider et al., 2011), with each substitution assumed to have resulted from a distinct mutational event. However, the distribution of SNPs across the genome has been known for some time to not be even, with not only polymorphisms (Amos, 2010; Hodgkinson and Eyre-Walker, 2010) but also fixed differences between species clustering in the genome (Bazykin et al., 2004). Where multiple base substitutions are found in the same genomic region, the derived alleles are more often found on the same haplotype (Schrider et al., 2011), suggesting that the two changes have not occurred independently.

Directly neighbouring polymorphisms are particularly enriched in the human genome (Hodgkinson and EyreWalker, 2010), though why this is the case is not fully understood. Previous studies of trios have shown that many of these changes have arisen in a single generation as multinucleotide polymorphisms (MNPs) (Besenbacher et al., 2016; Schrider et al., 2011), which are particularly enriched with GA → TT and GC → AA changes, suggesting that they are linked to error prone replication by polymerase zeta (Harris and Nielsen, 2014). However, not all sites of neighbouring changes can be explained by a single mutational event. Many neighbouring polymorphisms occur at different allele frequencies in the population suggesting they have arisen from two distinct mutations (Hodgkinson and Eyre-Walker, 2010), which we term here sequential dinucleotide mutations (SDMs). These clustered changes have been comparatively understudied and may simply reflect mutational hotspots but may also be the result of selection, with the impact of an initial deleterious SNP being at least partly corrected by a second neighbouring change (Davis et al., 2009). One obvious circumstance is where the impact of a deleterious nonsynonymous change is offset by a second polymorphism nearby. For example, nonsynonymous changes are less often found on highly expressed haplotypes, suggesting selection has favoured particular combinations of coding and regulatory alleles (Lappalainen et al., 2011). Likewise there is evidence that selection has favoured particular combinations of nonsynonymous changes spanning different amino acids in the same protein (Breen et al., 2012).

In this study we therefore focused on SDMs comprising two neighbouring changes that cannot be readily explained by a single mutational event, and investigate whether they simply reflect two independent but neighbouring polymorphisms or whether the second change depends on the first. The human mutation spectrum has diverged between human populations, with particular SNPs in particular sequence contexts more common in different continental groups (Harris and Pritchard, 2017). Accordingly, we also explore whether SDM fractions have diverged between population groups and whether any divergence simply mirrors differences in SNP or MNP mutational profiles. We also characterise the functional importance of these SDMs by studying the selective pressures acting upon them and whether they have favoured particular pathways through the amino acid code.

## Results

To investigate the frequency of occurrence of neighbouring mutations in different populations we defined SDMs in the 1000 genomes phase 3 dataset (The 1000 Genomes Project Consortium, 2015) using the approach illustrated in Fig 1. In this study we focused specifically on changes across neighbouring nucleotides that, due to being found at different allele frequencies, cannot be readily explained by a single mutational event. All of these SDMs were annotated with respect to their ancestral alleles, triplet nucleotide context and occurrence in each individual (see methods), allowing us to infer the order of nucleotide changes. SNPs were defined in the same way to enable direct comparisons of their mutational profiles to SDMs.

**Figure 1.**
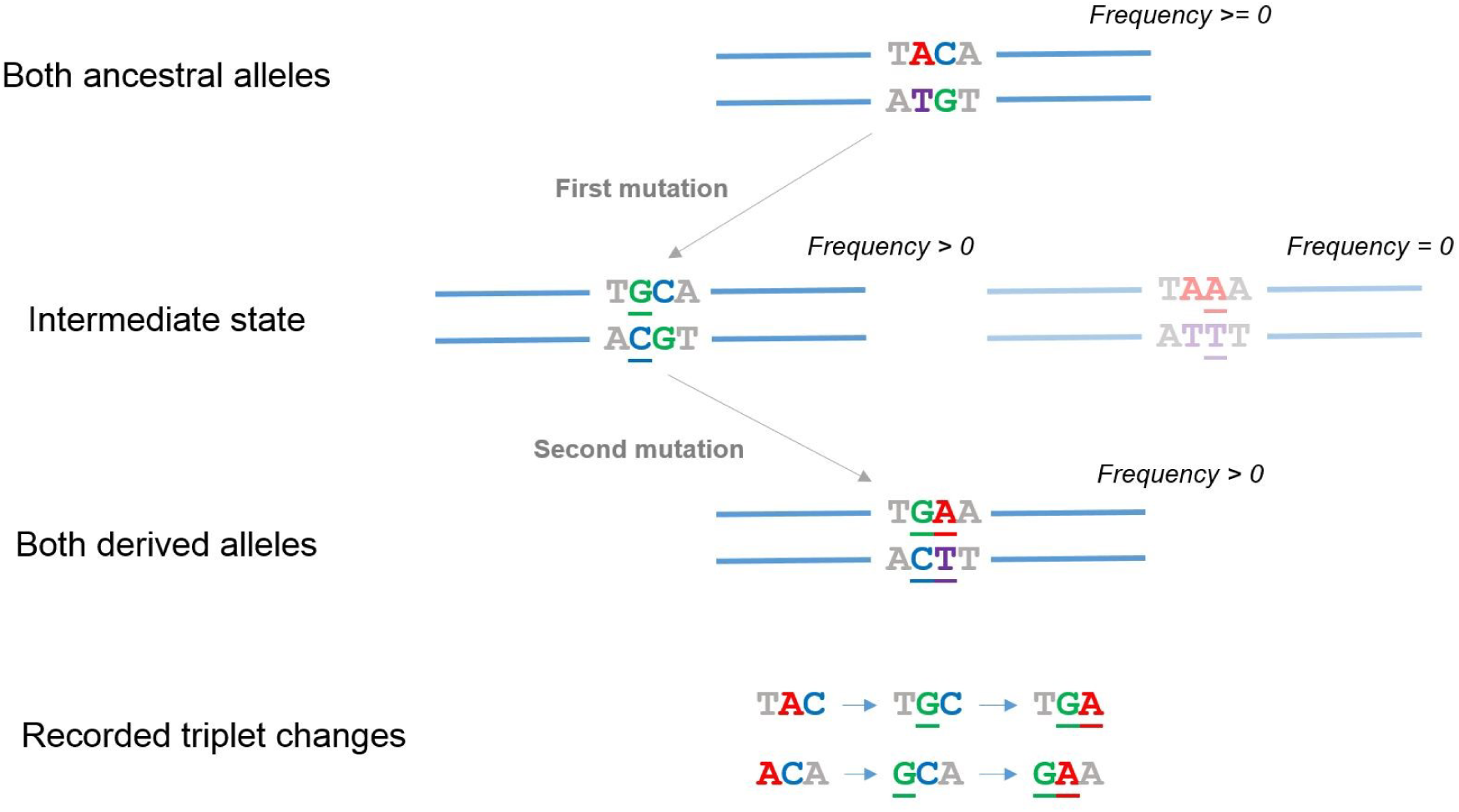
Defining the SDMs studied in this analysis. In this study we focused on SDMs involving neighbouring nucleotides that could not be readily explained by a single mutational event. Haplotypes containing just one of the two changes that make up the SDM had to be observed among the individuals (we assumed that reverting mutations and recombination events between neighbouring bases were rare). Using information on which alleles were ancestral we then inferred the order of changes, with each SDM recorded twice, according to their immediate 5’ and 3’ nucleotides. This allowed downstream analysis of their impact on codons and comparisons to SNPs in the same triplet contexts.

We observed that a typical individual carries over 14,000 of these compound SDMs (Supplementary Figure S1) of which on average 27 fall within a protein coding region. The distribution of SDMs across the genome broadly follows that of SNPs (Supplementary Figure S2 to Supplementary Figure S6), with the exception of the major histocompatibility complex on chromosomes 6, that carries an unusually high proportion of SDMs despite its high SNP density (Supplementary Figure S7). Examination across populations highlights that the intermediate haplotype is generally common (Supplementary Figure S8) and for 88% of SDMs it is observed across all continental groups, suggesting that the first change for many SDMs occurred prior to the human migration out of Africa (Fig. 2A).

**Figure 2.**
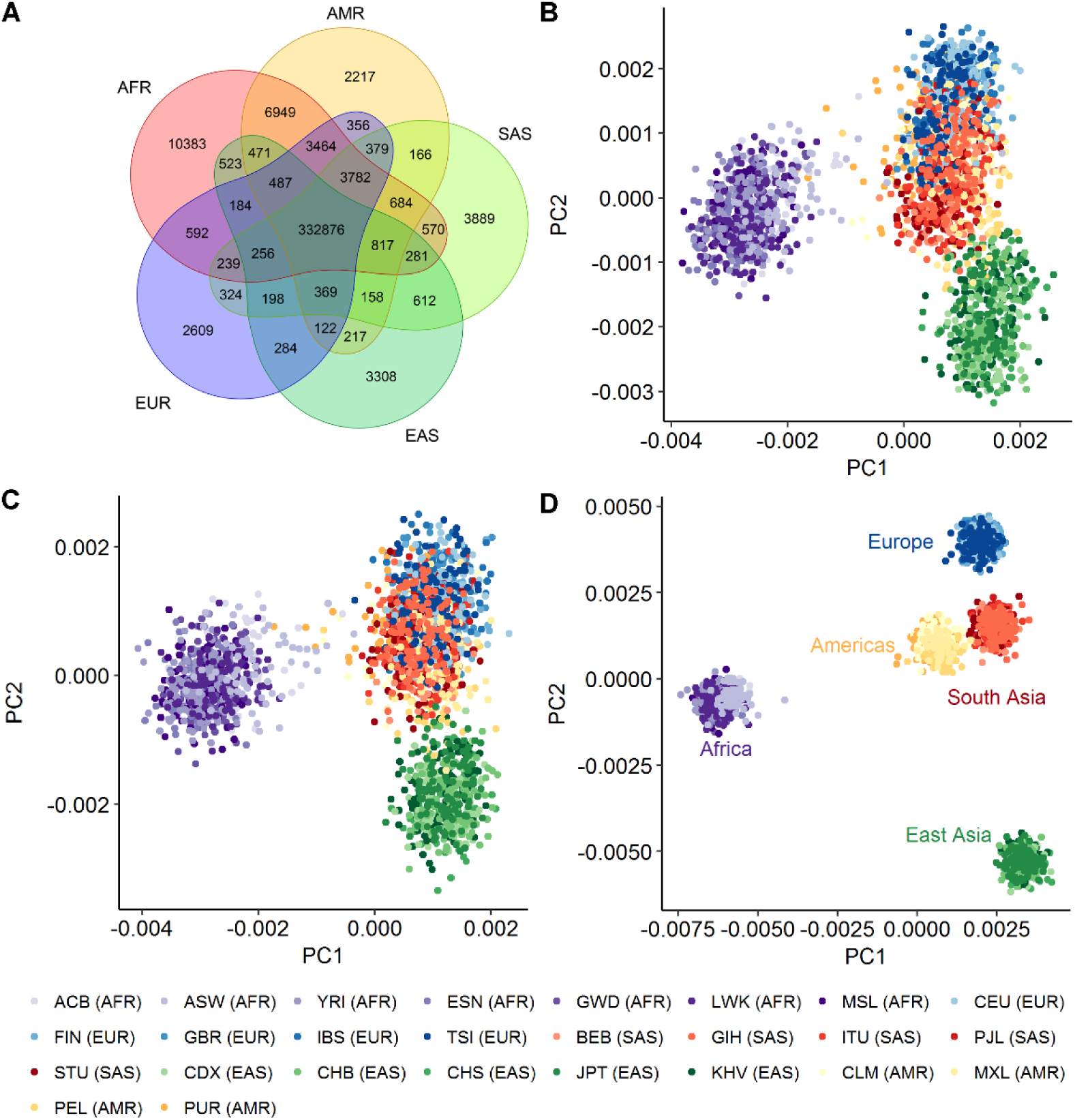
Divergence in SDM mutational fractions between human populations. A) Venn diagram of the number of intermediate haplotypes observed across different populations. B) Principal component analysis of individuals according to the mutational fraction of each first change in the SDMs they carry, defined by their triplet context. Points are coloured according to the continental group to which they belong and the corresponding 1000 genomes consortium three letter population codes are shown below. Although restricted to SNPs forming part of SDMs only, this plot largely mirrors that derived using all SNPs in the same populations presented in Harris and Pritchard (2017) at https://doi.org/10.7554/eLife.24284.006.C) The same as A but for the second change in SDMs. D) Principal component analysis of individuals according to the mutational fraction of the SDM changes they carry defined by triplet context.

### SDMs show biases between human populations distinct to those among SNPs

Despite being a reduced summary statistic compared to individual genotypes, comparisons of the numbers of SNPs found in different triplet contexts between individuals has been shown to separate out the major human population groups (Harris and Pritchard, 2017). This has been attributed to differences in the large number of genes that control DNA mutation and repair between populations. Characterisation of the individual SNPs that make up SDMs recapitulates the patterns observed in this previous study. Principal component analysis of the frequency of occurrence of the first and second changes in compound SDMs, defined by their sequence context, show highly similar patterns to that observed in Harris and Pritchard (2017) (Fig 2B and C). Despite their more limited numbers, the mutational profile of SDMs can though even more effectively separate out the major human sub-populations (Fig 2D); with certain types of SDMs in specific nucleotide contexts enriched in different populations. Surprisingly, unlike the PCAs of individual SNPs, the American continental group separates in this analysis, in part driven by a depletion for CG → TG → TA SDMs among these individuals (Fig 3A). In contrast SDMs are relatively depleted at AT rich triplets in African populations, and in particular SDMs where the mutations have the effect of switching AT base composition between strands (i.e. that involve multiple neighbouring A:T → T:A or T:A → A:T mutations; Fig 3B). To ensure these results were not due to ancestral allele misidentification we repeated the analysis but this time without restricting to sites where the ancestral allele could be determined. After grouping sites sharing the same combinations of haplotypes, the major populations were still observed to separate in this analysis (Supplementary Figure S9). A PCA based on MNP mutational fractions, i.e. those changes for which an intermediate haplotype is not observed, also does not show the definition between continental groups observed for SDMs (Supplementary Figure S10).

**Figure 3.**
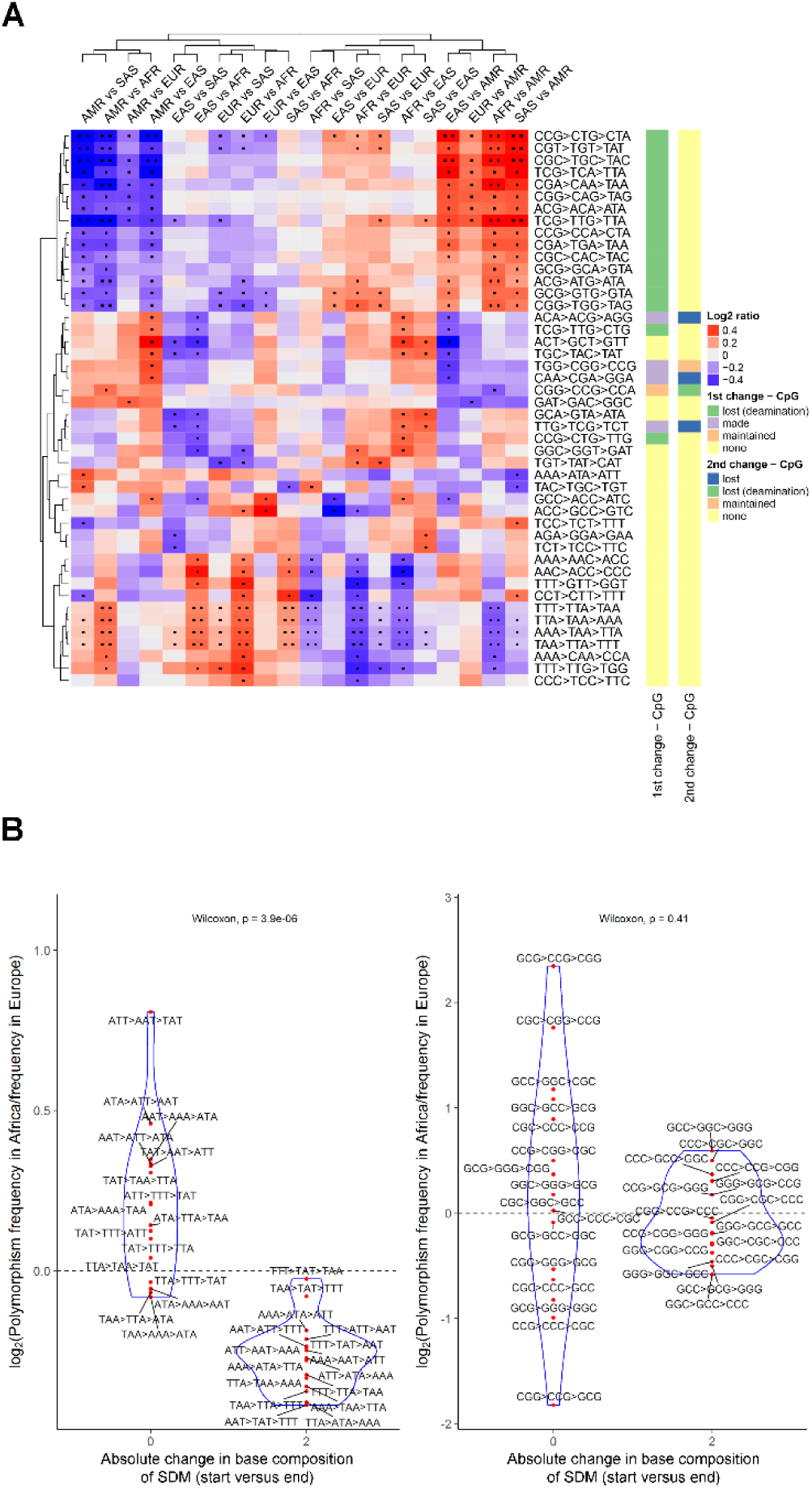
SDMs that differ between population groups. A) Log^2^ ratios of the frequencies with which selected SDMs occur in different continental groups. Only SDMs found at at least 150 sites in each population and with a Log_2_ratio >= 0.3 and a P < 0.05 in at least one population comparison are shown in this plot. One dot indicates the corresponding comparison was associated with an uncorrected Fisher’s exact P < 0.05, two dots that the false discovery rate (q value) was < 0.05. Red cells indicate the corresponding SDM is relatively enriched in the first named continental group, blue that the change is enriched in the second population. B) The relative enrichment of selected SDMs in African versus European populations. (Left) SDMs that exclusively involve weak base pairs (adenine and thymine) broken down by their net impact on the base composition of each strand. (Right) SDMs that exclusively involve strong base pairs (guanine and cytosine).

This improved separation observed in the SDMs analysis is driven by the fact that the first and second changes in SDMs show distinct biases between populations, providing a greater resolution of population differences. This is illustrated by the fact a PCA of the differences in mutational fractions between the first and second change in SDMs also effectively separates the major population groups (Supplementary Figure S11). If the first and second change in SDMs showed the same biases between populations, then continental groups should not still separate in this analysis.

Consequently, SDMs show different mutational spectra across populations that are distinct and more pronounced than those among SNPs and MNPs, and which can effectively separate major population groups.

### CpG deamination is not sufficient to explain SDM variation

Intriguingly, the results in Supplementary Figure S11 imply that the first and second mutations in SDMs are not driven by the same processes. To investigate this further we characterised the types of mutations observed as the first and second change in SDMs classified again by base change and triplet context. To ensure that batch effects and sequencing artefacts did not confound this analysis we replicated it across two independent datasets. The global set of 2504 genomes sequenced by the 1000 genomes consortium used above, as well as a novel dataset of 1358 Scottish genomes from the Lothian Birth Cohorts 1921 and 1936 (Deary et al., 2012; Taylor et al., 2018), whole genome sequenced at a mean depth of 36X. Comparison of the frequency with which particular changes occur as the first and second mutation in an SDM indicates that each shows biases for different types of change. As shown in Fig 4 a clear role of CpG dynamics in shaping SDMs is observed. Methylated cytosines immediately followed by a guanine (i.e. CpG sites) are known to be particularly prone to deaminate to a thymine, with mutation rates at these sites up to 18 times higher than at other dinucleotides (Kong et al., 2012). As shown in Fig 4, the first mutation in SDMs is more likely to create a new CpG site, with the second change more likely to lead to the loss of one. This suggests that a dominant factor underlying SDMs is an initial mutation that creates a new CpG site, which subsequently mutates. This signature is observed across both the 1000 genomes and Lothian Birth Cohorts (Supplementary Figure S12).

**Figure 4.**
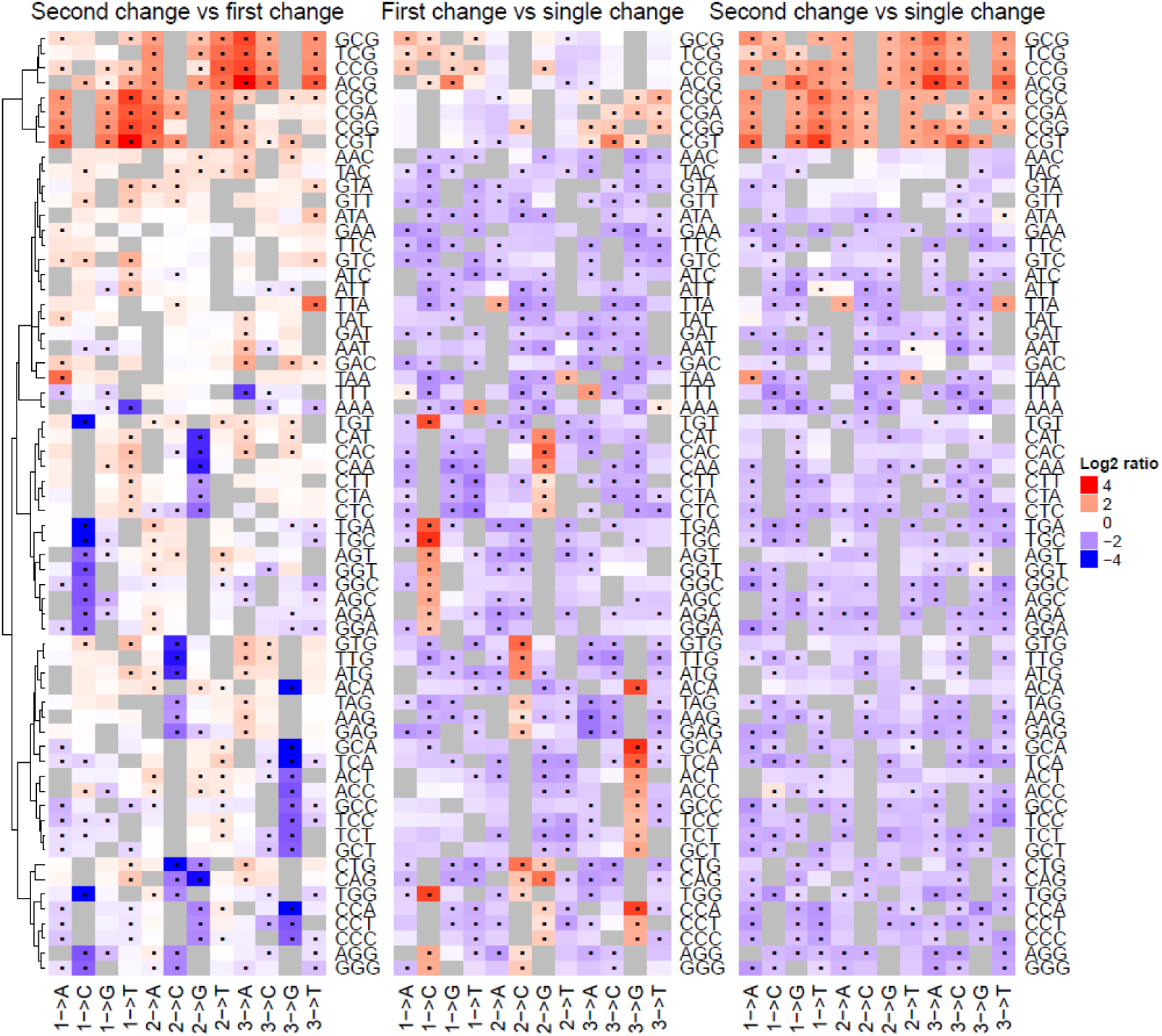
The relative mutational fractions of different forms of first and second changes among SDMs. (Left) The ratios of the mutation fractions of the first and second changes by triplet context in the 1000 genomes cohort. Each cell corresponds to a particular change defined by the original triplet (labelled on the right of the plot) and the observed change (labelled at the bottom). The numbers in the bottom row indicate which base in the triplet is polymorphic followed by the mutation that occurred i.e. 1-> A in the bottom GGG row indicates the respective cell corresponds to GGG-> AGG changes. A dot in the cell indicates that the frequency of the corresponding change is significantly different between the first and second changes of SDMs (Bonferroni corrected chi-squared P < 0.05, see methods for more details). (Middle and right) The ratio of the frequencies of the first (middle) and second changes (right) in SDMs versus the observed frequencies of the same change at SNPs. In all three heatmaps red cells indicate the change is enriched among the first named change, and blue that it is enriched among the second named change.

This raises the question, to what extent do SDMs simply reflect the known mutational biases of SNPs? (Supplementary Figure S13). To explore this we determined the expected number of each SDM in the genome given the observed frequencies of occurrence of its constituent changes among SNPs (see methods for more details). Moderate correlations were observed between these values (Fig 5A, negative bionomial regression McFadden’s psuedo-R^2^: 0.57 (Lothian Birth Cohort), 0.61 (1000 Genomes Cohort)), but substantial outliers were observed, where the frequencies of occurrence of SDMs in the genome differ markedly from what would be expected given the rates of changes among SNPs. In particular the SDMs involving CA → CG → TG changes (and their complement TG → CG → CA) occur over an order of magnitude more frequently than expected given the frequency of occurrence of the constituent changes among SNPs (Fig 5A). Notably, they also occur over an order of magnitude more often than other changes that involve the creation and subsequent deamination of CpG sites. Whereas 10,633 intergenic CAG → CGG → TGG changes were observed in the 1000 genomes population there are only 1,038 intergenic CTG → CGG → TGG changes, despite both changes leading to a similar creation and loss of CpG sites, and both ancestral triplets occurring at the same frequency in the genome (due to being the reverse complement of one another). This enrichment of these changes is maintained at extended sequence contexts (Supplementary Figure S14). This implies that a distinct process where an initial A:T → G:C change is favoured has led to the comparatively high number of these changes, and CpG dynamics alone cannot explain the elevated occurrence of these specific SDMs relative to other SDMs involving a final CpG to TpG change. This is further illustrated in Supplementary Figure S15. Modelling the interaction between these two factors (original base change and the impact of changes on CpG sites) using regression analysis confirms that only where a CpG is created by an initial A → G, and not for example by a T → G change, is the rate of SDMs so high (Supplementary Figure S15).

**Figure 5.**
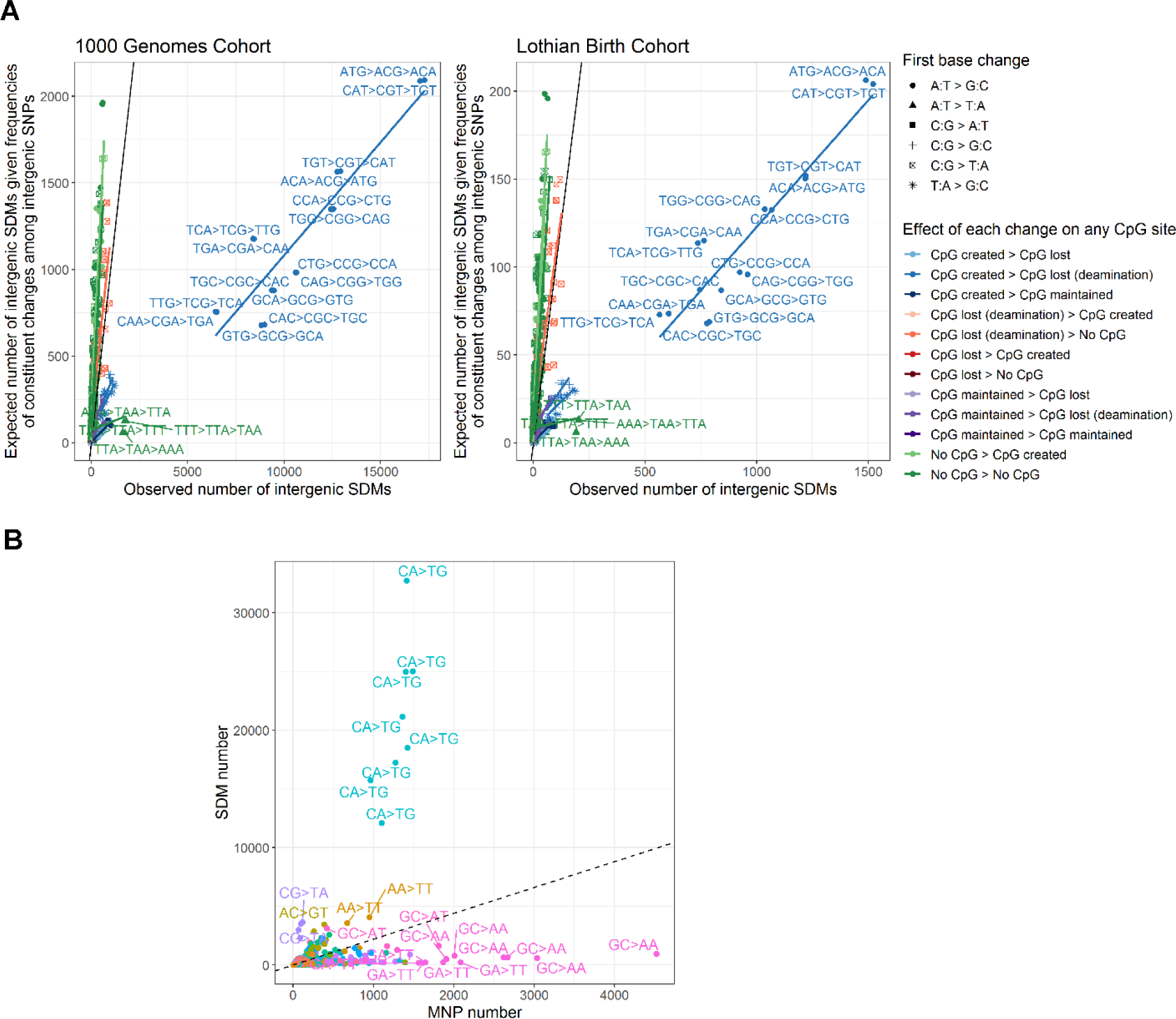
The observed number of intergenic SDMs against the number expected given the frequencies with which the constituent changes occur among SNPs and MNPs. A) SDMs versus SNPs. SDMs are broken down by the original base change and the impact on any CpG sites of each constituent change (impact of first change -> impact of second change). The line of parity is shown in black. Together these three factors (the expected number of SDMs from background SNP rates, the first base change and impact of the changes on CpG sites) explain the majority of the variation in SDM frequencies (McFadden’s pseudo-R^2^: 0.84 (Lothian Birth Cohort), 0.85 (1000 Genomes Cohort)). B) The number of SDMs versus number of MNPs showing the same ancestral and derived triplets in the 1000 genomes cohort. Points are coloured by the dinucleotide change between the ancestral and derived haplotypes and outlier points labelled.

Comparison of the number of SDMs to the number of MNPs sharing the same ancestral and derived triplets in the 1000 genomes cohort further highlights the distinct mutational biases of SDMs and in particular the bias towards the specific enrichment for CA → CG → TG changes (Figure 5B). Whereas, as previously observed, MNPs are enriched with GA→TT and GC→AA changes, thought to be a result of error prone replication by polymerase zeta (Harris and Nielsen, 2014), SDMs are distinct due to this bias towards CA → CG → TG mutations.

Further mutational biases specific to SDMs are also observed. For example TTA to TTT polymorphisms are more common as the second change in an SDM than the first, and SDMs containing two consecutive A:T→ T:A changes are more common than expected given the frequency of occurrence of the same changes among SNPs (Fig 4 and Fig 5A). Consequently although the turnover of CpG sites drives the creation of a large proportion of SDMs, further processes appear to be contributing to different biases not only between SDMs and SNPs/MNPs, but also between the first and second changes of neighbouring polymorphisms.

### The sequence of changes observed depends on their ancestral state

This relationship between the first mutation in SDMs and their frequency of occurrence in the genome raises the question as to whether the second mutation depends to some extent on the first. To explore this, we examined SDMs where initial mutations have created the same intermediate nucleotide triplet. If the two changes that comprise an SDM are independent, then the subsequent mutations at these intermediate triplets should occur in similar numbers irrespective of the ancestral triplet from which they derived.

The dominant feature underlying downstream changes at many of these intermediate triplets was the biased deamination of CpG sites. SDMs passing through an intermediate triplet containing a CpG site are dominated by subsequent CpG to TpG changes irrespective of the original ancestral nucleotides. We therefore focused on intermediate triplets that contain no CpG sites. As shown in Fig 6A, mutations at these intermediate triplets are not independent of the initial change. For example, 94% of SDMs passing through an intermediate TTA triplet go on to become TAA if the original ancestral codon was TTT. This number is though only 17% if the original ancestral codon was TTG (Fig 6A and Table 1). This link between the form of the first change on the observed fraction of downstream changes was shown to be maintained in expanded sequence contexts (one and two bases either side of each SDM, see Supplementary Table 1). To minimise the potential impact of batch effects and sequencing artefacts we again sought replication for this observation in the independent Lothian Birth Cohort collection of Scottish genomes and this difference was found to be highly significant in both datasets (Fig 6B). This phenomenon is not exclusively restricted to intermediate triplets containing just adenines and thymines. For example, ACA intermediate triplets are more likely to be linked to an ATA final triplet if the ancestral triplet was CCA than if it was ACG (Fig 6A+B). There appear therefore to be constraints on the second change in SDMs dependent on their original ancestral state.

**Figure 6.**
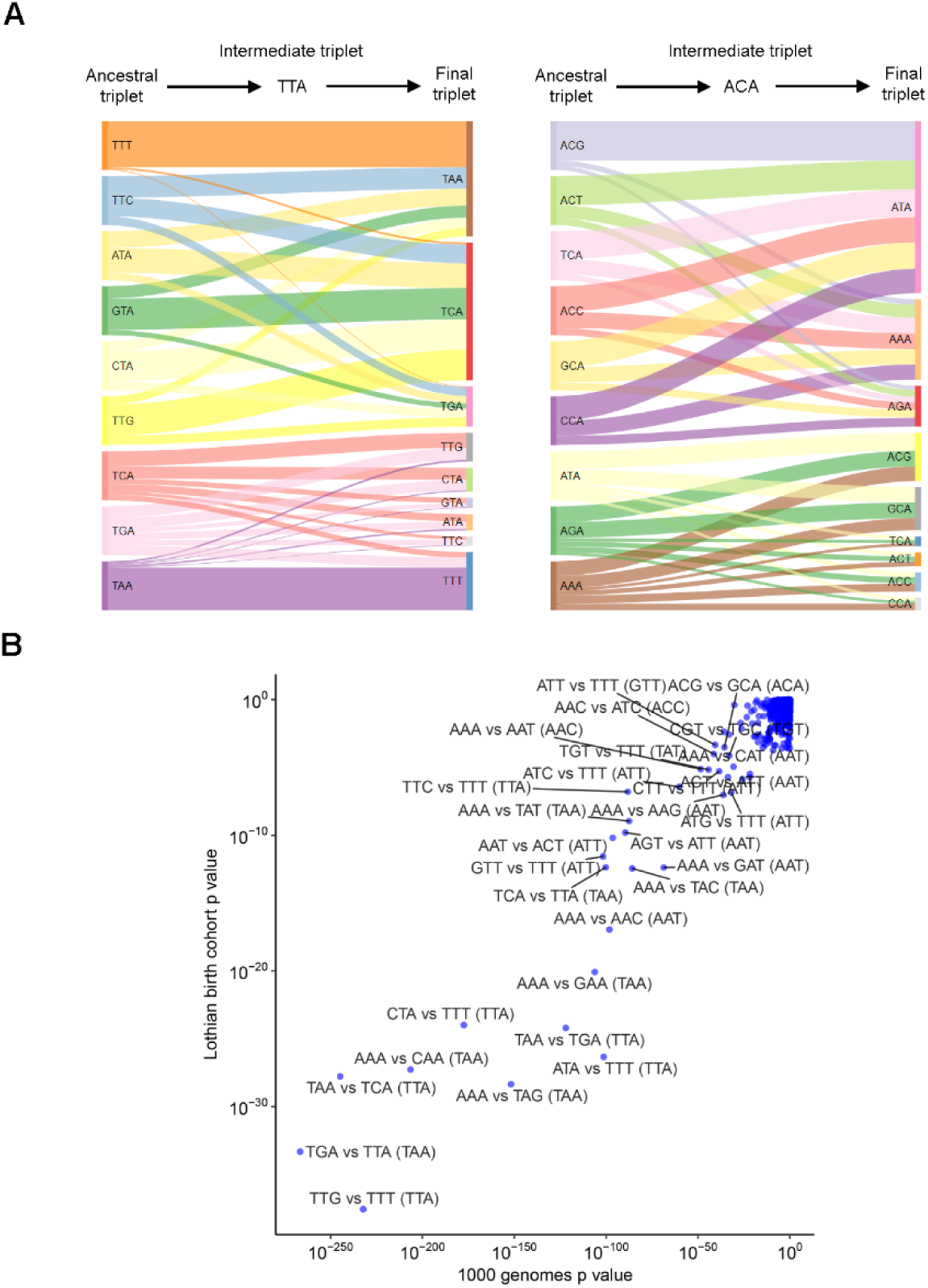
The form of the second change is related to the ancestral state of SDMs. A) Sankey plots representing the changes observed among intergenic SDMs sharing the same intermediate state. The width of each bar represents the proportion of SDMs sharing the corresponding ancestral state that ended at the indicated final triplet. B) P values indicating pairs of ancestral states where intergenic SDMs show different biases for final triplets despite passing through the same intermediate state (indicated in brackets). Chi-squared test p values for both the 1000 genomes and Lothian Birth Cohorts are shown.

**Table 1.** Counts and proportions (by ancestral triplet) of SDMs passing through the intermediate triplet TTA.

| Ancestral triplet | Intermediate triplet | Derived triplet | Intergenic SDMs | Intergenic proportion by start triplet | Intronic SDMs | Intronic proportion by start triplet | Coding SNP count (first change) | Coding SNP count (second change) |
| --- | --- | --- | --- | --- | --- | --- | --- | --- |
| ATA | TTA | TAA | 38 | 0.34 | 32 | 0.42 | 314 | 68 |
| ATA | TTA | TCA | 56 | 0.50 | 36 | 0.47 | 314 | 558 |
| ATA | TTA | TGA | 17 | 0.15 | 9 | 0.12 | 314 | 91 |
| CTA | TTA | TAA | 36 | 0.21 | 20 | 0.19 | 1207 | 68 |
| CTA | TTA | TCA | 104 | 0.61 | 69 | 0.64 | 1207 | 558 |
| CTA | TTA | TGA | 30 | 0.18 | 18 | 0.17 | 1207 | 91 |
| GTA | TTA | TAA | 21 | 0.25 | 19 | 0.30 | 403 | 68 |
| GTA | TTA | TCA | 54 | 0.64 | 34 | 0.53 | 403 | 558 |
| GTA | TTA | TGA | 9 | 0.11 | 11 | 0.17 | 403 | 91 |
| TTC | TTA | TAA | 62 | 0.44 | 55 | 0.49 | 656 | 68 |
| TTC | TTA | TCA | 54 | 0.38 | 43 | 0.38 | 656 | 558 |
| TTC | TTA | TGA | 25 | 0.18 | 15 | 0.13 | 656 | 91 |
| TTG | TTA | TAA | 44 | 0.17 | 38 | 0.20 | 1865 | 68 |
| TTG | TTA | TCA | 162 | 0.64 | 121 | 0.64 | 1865 | 558 |
| TTG | TTA | TGA | 49 | 0.19 | 31 | 0.16 | 1865 | 91 |
| TTT | TTA | TAA | 1817 | 0.94 | 1696 | 0.95 | 257 | 68 |
| TTT | TTA | TCA | 81 | 0.04 | 55 | 0.03 | 257 | 558 |
| TTT | TTA | TGA | 28 | 0.01 | 27 | 0.02 | 257 | 91 |
| TAA | TTA | ATA | 61 | 0.03 | 52 | 0.03 | 15 | 168 |
| TAA | TTA | CTA | 46 | 0.02 | 44 | 0.03 | 15 | 606 |
| TAA | TTA | GTA | 27 | 0.01 | 19 | 0.01 | 15 | 244 |
| TAA | TTA | TTC | 23 | 0.01 | 18 | 0.01 | 15 | 182 |
| TAA | TTA | TTG | 76 | 0.04 | 47 | 0.03 | 15 | 1046 |
| TAA | TTA | TTT | 1611 | 0.87 | 1489 | 0.89 | 15 | 209 |
| TCA | TTA | ATA | 80 | 0.15 | 55 | 0.14 | 1503 | 168 |
| TCA | TTA | CTA | 130 | 0.25 | 90 | 0.23 | 1503 | 606 |
| TCA | TTA | GTA | 58 | 0.11 | 43 | 0.11 | 1503 | 244 |
| TCA | TTA | TTC | 35 | 0.07 | 34 | 0.09 | 1503 | 182 |
| TCA | TTA | TTG | 160 | 0.31 | 127 | 0.33 | 1503 | 1046 |
| TCA | TTA | TTT | 60 | 0.11 | 39 | 0.10 | 1503 | 209 |
| TGA | TTA | ATA | 32 | 0.14 | 11 | 0.06 | 24 | 168 |
| TGA | TTA | CTA | 47 | 0.21 | 29 | 0.17 | 24 | 606 |
| TGA | TTA | GTA | 19 | 0.09 | 16 | 0.09 | 24 | 244 |
| TGA | TTA | TTC | 25 | 0.11 | 18 | 0.10 | 24 | 182 |
| TGA | TTA | TTG | 54 | 0.24 | 43 | 0.25 | 24 | 1046 |
| TGA | TTA | TTT | 45 | 0.20 | 55 | 0.32 | 24 | 209 |

### Coding SDMs have favoured the creation of single codon amino acids

We next investigated the impact of these mutational biases on coding regions. As shown in Fig 7, SDMs have had a distinct impact on the evolution of genes. The particular biases of SDMs means that nine out of ten of the most common coding SDMs involve the previously described CA → CG → TG change. As a result of the layout of the amino acid code this bias has led to the preferential creation of the single codons that code for methioinine and tryptophan (logistic regression of frequencies of changes that create these single codon amino acids to the frequencies of all other changes: P = 6.4×10^−04^). Thr_ACA_ → Thr_ACG_ → Met_ATG_ is the most common coding SDM, with Gln_CAG_ → Arg_CGG_ → Trp_TGG_ the fourth most common change (Fig 7, Supplementary Figure S16). The mutational biases of SDMs and organisation of the amino acid code have therefore combined to favour the creation of single codon amino acids. The comparatively low number of these amino acids created by SNPs is therefore partially compensated for by the particular mutational biases of SDMs.

**Figure 7.**
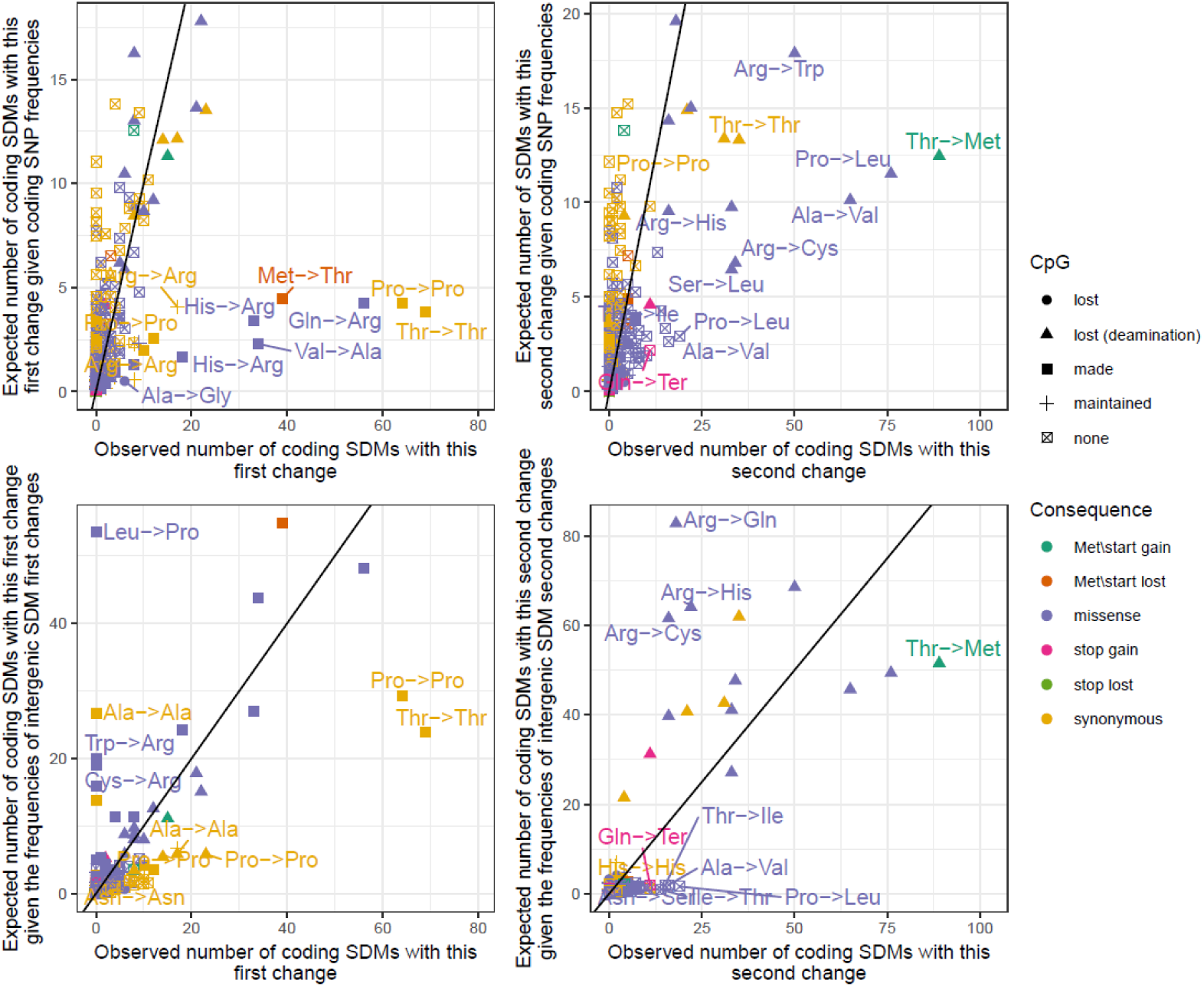
The observed and expected number of coding SDMs in the 1000 genomes cohort. The x axes in the left and right columns correspond to the observed number of first and second changes in coding SDMs respectively. Each point represents a distinct single base change between two codons and points are coloured by the impact of the change on the corresponding protein. The impact of the change on any CpG sites is also indicated by the shape of the point. The y axis in the top row corresponds to the number of the changes expected given their frequency among coding SNPs. The y axis in the bottom row gives the expected numbers given the frequencies of the changes amongst the first and second changes of intergenic SDMs after accounting for differences in the frequencies of occurrence of ancestral triplet between regions. Labelled points show a significant difference between the observed and expected numbers after correcting for multiple testing (Chi-squared P< 2.4e^-5^). The line of equality is also indicated.

To investigate the potential impact of selection in coding regions we compared the observed number of coding SDMs to that expected given the numbers of the same SDM in intergenic regions, after accounting for differences in the occurrence of the ancestral triplets between regions (see methods). Under the assumption that intergenic SDMs are under comparatively little selection, discrepancies between these numbers should indicate which coding SDMs have been favoured or disfavoured by selection. A noticeable difference to the previous comparison with coding SNPs is that the first change of coding SDMs is depleted with a number of the changes that create a CpG site (Fig 7, Supplementary Figure S17). SDMs where the first mutation is non-synonymous (missense, stop gained or lost) are depleted among coding regions, consistent with their removal by selection, with the notable exception of the Thr_ACA_ → Thr_ACG_ → Met_ATG_ pathway through the amino acid code that remains unusually enriched among coding SDMs (observed proportion of coding SDMs matching this change versus expected proportion given number in intergenic regions and differences in rates of ancestral codon between regions: Chi-squared P=2.9×10^-10^, Bonferroni corrected P = 6.58×10^-7^). This suggests this change is not only favoured by the mutational biases of SDMs but also by selection in coding regions.

To explore the mutational profiles of coding SDMs further, we compared the normalised occurrence with which SDMs occur on the coding and non-coding strands of genes. Taking the SDM shown in Fig 8A as an example, as a change on one strand must also effect the other strand then the null hypothesis is that the numbers of these two complementary changes (CGA>GCG>GTG and TGC>CGC>CAC) should be similar. In this analysis we compared if though there was an impact of whether a gene was present on the blue or red strand. If the frequencies of these changes are independent of whether or not they occur on the coding strand then the two changes should still occur at the same rates when accounting for the background occurrence of the respective ancestral codon on the coding strand of genes. This is what we see for most changes, i.e. an SDM and its reverse complement are found at approximately equal normalised occurrences on the coding strand of genes (Fig 8B). However, a subset of SDMs show imbalances in the normalised occurrence with which they and their reverse complement occur on the coding strand, including both the Thr_ACA_ → Thr_ACG_ → Met_ATG_ and Gln_CAG_ → Arg_CGG_ → Trp_TGG_ changes. Of the five most significant SDMs in this analysis, only one; Pro_CCG_ → Pro_CCA_ → Leu_CTA_ does not involve a CA → CG → TG change. The reverse complement of this change is ArgCGG → TrpTGG → Stop_TAG_ suggesting that this change is comparatively infrequent on the coding strand due to it leading to the introduction of a deleterious stop codon. Consequently selection acting on a subset of SDMs appears to have led to imbalances with which they and their reverse complement appear on the coding strand of genes.

**Figure 8.**
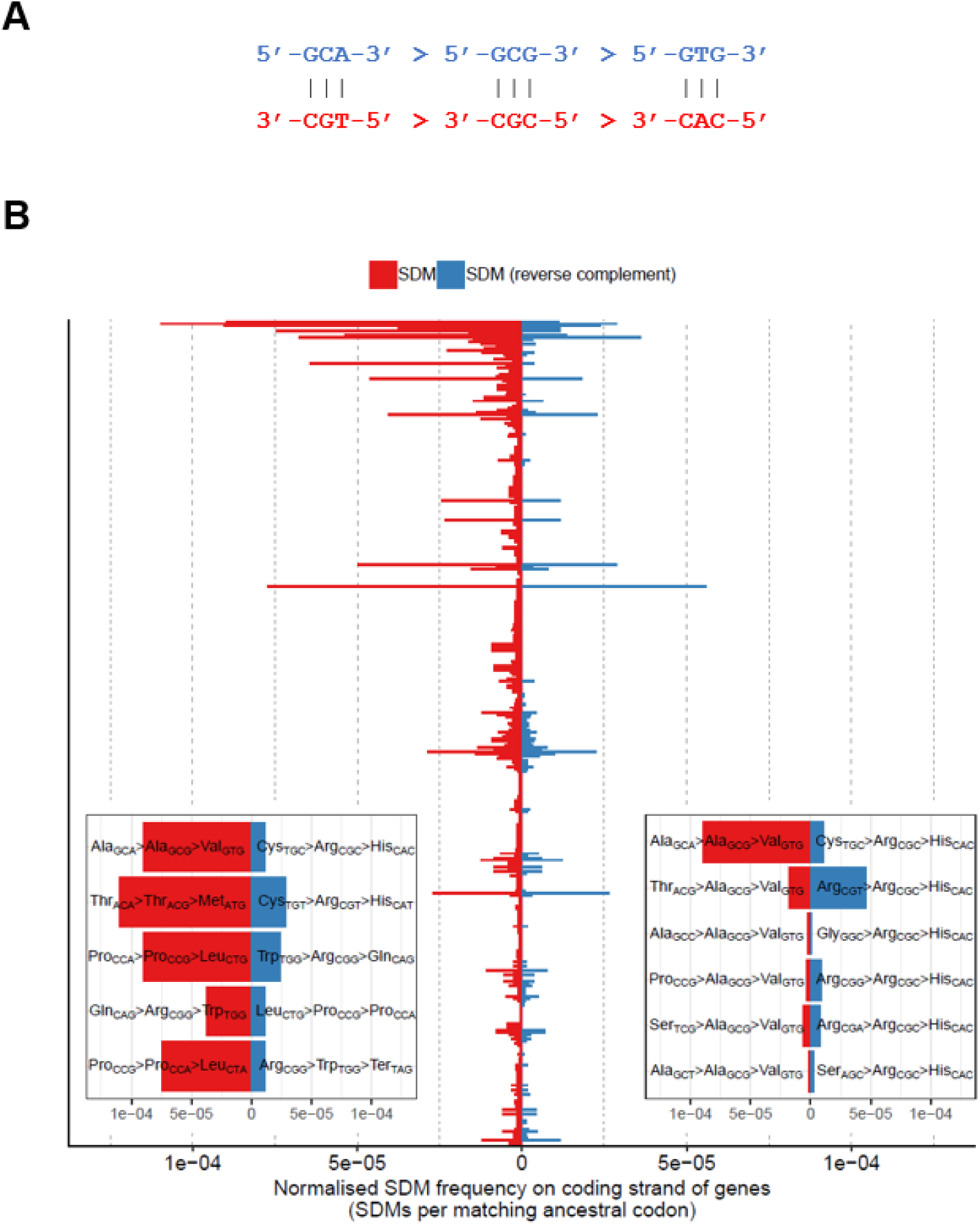
Asymmetry in the occurrence of changes on the coding and non-coding strand of genes. A) Reverse complement changes are expected to occur at the same rates across the genome. B) The observed count of each SDM change on the coding strand of genes, normalised by the total frequency with which the ancestral codon occurs in coding regions, is shown. Each change is paired with its reverse complement change, with the count of the change displaying a comparative enrichment on the coding strand shown in red. Where both changes are equally common on coding strands, one was randomly chosen to be shown in red. To test for asymmetry in the frequencies of SDM pairs, a 2×2 contingency table of (a) the count of each SDM in coding regions and (b) the count of the matching ancestral codons across all genes was constructed, and differences in the proportion of ancestral codons carrying the respective SDM was tested using the Fishers exact test. SDM pairs are ranked by p value (smallest at the top). The five most significant pairs (P <5×10^-5^) are shown in the bottom-left inset and the six SDM involving a final Ala_GCG_ -> Val_GTG_ change are shown in the bottom right.

We conclude that although particular changes are favoured by SDM mutational biases, selection has preferentially removed changes passing through certain codons, leaving changes that create single codon amino acids comparatively unaffected. Together, this has led to different impacts on coding regions of SDMs and SNPs.

### Evidence for epistatic selection at neighbouring coding polymorphisms

We finally investigated whether there is evidence of epistatic selection acting across the polymorphisms that make up coding SDMs i.e. whether the selective pressure acting on a nucleotide change depends on neighbouring changes. The amino acid code is thought to have been optimised so that physically similar amino acids have been brought together at neighbouring positions (Koonin and Novozhilov, 2009). This ensures that the negative effect of a single mutation is minimised. By extension though, this suggests that two successive missense mutations in a codon are potentially more deleterious than one, despite the net change being one amino acid change in both cases, with the strength of selection acting on the first mutation modulated by the allele present at the neighbouring base.

To explore this hypothesis we focused on SDMs where the initial mutation led to a missense change, and characterised whether the strength of selection acting upon this initial change depends upon subsequent neighbouring mutations in the same codon. If a subsequent change in the same codon is synonymous then the original amino acid change is unaffected and the null hypothesis is that the strength of selection on the codon should be in line with that among coding SNPs with the same change as the original missense mutation. However, if the second change is a further non-synonymous change and the amino acid code is optimised so that two subsequent missense changes in the same codon are more deleterious than just one, such changes should be relatively depleted in the genome despite the fact there is still only one amino acid change. We used multiple linear regression to control for potential confounders (the occurrence of the first change among coding SNPs indicating the expected occurrence of these changes, the number of different codons that encode the final amino acid and the impact on any CpG site of the second change). Supplementary Figure S18 shows that SDMs made up of two successive missense changes are less frequently observed in the genome when compared to missense changes followed by a subsequent synonymous change (P=0.0013, false discovery rate=0.0088). The number of missense-synonymous SDMs in the genome is on average 37% higher than missense-missense SDMs after accounting for the number of coding SNPs showing the same change as the original missense mutation, CpG mutability and codon frequency. When also accounting for the state of the intermediate triplet, this difference remains significant (P=0.00015, false discovery rate=0.010).

Although single codon amino acids appear to have been generally favoured by SDMs, having accounted for this effect, the creation of a methionine codon following an original missense change in fact occurs particularly infrequently relative to synonymous changes (P=1.4×10^-6^, false discovery rate=1.3×10^-5^). Consequently the impact of missense changes on fitness appears to depend on the form of subsequent neighbouring mutations, and epistatic selection between neighbouring coding polymorphisms has helped shape the number of SDMs in the human genome.

## Discussion

Although the changes that make up SDMs have been most often categorised as individual SNPs, we have shown they appear to be driven by distinct mutational processes. Although the creation and subsequent deamination of CpG sites underlies a large proportion of SDMs, we show, we believe for the first time, that CA → CG → TG changes are substantially overrepresented relative to other changes that involve the gain and deamination of a CpG site. This suggests a distinct process is driving this bias for these specific changes. The creation of new G:C base pairs is often attributed to biased gene conversion (BGC) that favours weak (A:T) to strong (G:C) basepair changes. However, previous studies have found little evidence of a strong effect of BGC on genome-wide mutational profiles (Harris and Pritchard, 2017; Do et al., 2015), and, for example, T:A → G:C changes are not similarly enriched among the first change of SDMs, suggesting a distinct process is potentially contributing to the elevated rate of these polymorphisms.

As with SNPs, SDM mutational profiles appear to differ between human groups, but, unlike SNPs and MNPs, SDM rates can more clearly define populations. In contrast to SNP mutational profiles, the SDM profiles of the Americas populations distinguishes them from other continental groups, in part driven by a depletion of CG → TG → TA changes among these individuals. The larger spectrum of changes among SDMs, and the fact that the changes in SDM are not independent, and do not simply reflect underlying SNP rates, likely provides the greater resolution in defining populations. Although genotyping and switch errors would impact the ability to call SDMs accurately, previous analyses have suggested that the accuracy of the 1000 genomes cohort is high and the use of the independent and high coverage (>30X) Lothian Birth Cohort to validate key findings suggests that errors due to low sequencing coverage in 1000 genomes samples aren’t driving the observations in this study. Our examination of switch errors in 26 randomly selected individuals through read based phasing suggests switch errors are relatively infrequent at these neighbouring bases. Likewise SDMs exhibit distinct mutational biases to MNPs suggesting they are not simply inappropriately annotated MNPs.

Not all common SDMs are associated with CpG sites. In particular, changes at AT rich triplets show population differences and enrichment among African populations. The second change in AT-rich SDMs also often appears dependent on the original ancestral sequence. One potential explanation for this is a role for homologous recombinational repair among these SDMs. Recombination based repair mechanisms transfer nucleotide sequence information between chromosome copies, and as a result between ancestral and derived haplotypes. The second changes in these SDMs may therefore reflect errors in this repair process, potentially arising due to the pre-existing mismatch between chromosomal copies. Alternatively constraints on the sequence composition of regions may lead to biases towards those maintaining local nucleotide composition. Recent work has highlighted that as many tests of adaptive evolution, such as the branch-site test, assume base substitutions occur independently, then violations of this assumption can lead to erroneous signals of positive selection (Venkat et al. 2018). MNMs have been show to drive a lot of false positive signals but this potentially also extends to SDMs.

An intriguing consequence of the bias for the creation and subsequent loss of CpG sites among SDMs is the fact that this favours the creation of methionine and tryptophan codons due to the specific arrangement of the amino acid code. As both codons contain a TG dinucleotide they are readily created by the preference for CA → CG → TG changes. This therefore partly offsets the fact that these amino acids are more rarely created by single mutations due to being only encoded by a single codon. Among coding SDMs the creation of new methionine codons is the most common of all double mutations, even when accounting for background mutation rates. However, it should be noted that as SDMs are relatively rare, the overwhelming majority of new methionine codons are still created by SNPs.

Nonadditive, i.e. epistatic, genetic interactions have been proposed to underlie a range of phenomenon such as the missing heritability of phenotypes, but detecting such interactions in humans has proven difficult (Wei et al., 2014). Previous studies have suggested that the strength of selection acting upon a coding variant may depend on the alleles carried at other variants nearby. For example Lappalainen et al. (2011) showed that putatively functional coding variants are less often observed on more highly expressed regulatory haplotypes. Given the previous observations that the amino acid code appears optimised so that the more deleterious amino acid changes cannot result from single base changes, we investigated whether SDMs may be a further example of epistatic selection in the human genome. When accounting for various factors, an original non-synonymous change is less often than expected followed by a non-synonymous change in the same codon. This suggests that the strength of selection acting upon the original missense mutation is modulated by changes next to it, despite the net effect still being a single amino acid change. However, an assumption made to varying degrees by these studies is that missense changes can be grouped, and that their impact on fitness is broadly similar. Even larger sequencing cohorts, such as those being generated as part of the UK Biobank (Sudlow et al., 2015), would help refine this analysis and the epistatic selection acting upon individual types of intermediate missense changes.

Consequently human mutation profiles appear more complex than previously thought, with neighbouring polymorphisms driven by distinct mutational processes. These SDMs are under unusually strong selective pressure and have played an important and distinct role in shaping human protein evolution.

## Materials and methods

### Variant calling

1000 genomes consortium version 3 phased haplotypes along with information on their ancestral alleles were obtained from http://ftp.1000genomes.ebi.ac.uk/vol1/ftp/release/20130502/. After excluding variants with missing or low confidence ancestral allele annotations, neighbouring variants where both derived alleles were observed together on the same haplotype were flagged as potential SDMs. As the focus of this analysis was on SDMs originating from two neighbouring mutation events, only SDMs where haplotypes also existed which carried the derived allele at one but not the other variant were kept, i.e. the two changes were observed at different allele frequencies. See Fig 1 for more details on how SDMs were defined in this study. This led to 169,702 putative MNPs being excluded where only two haplotypes were observed. Due to the very low probability of recombination events occurring between neighbouring bases in the human genome, we also excluded SDMs where both combinations of one derived and one ancestral allele were observed. Following this filtering 377,766 neighbouring pairs of polymorphisms remained.

As switch errors would impact the ability to correctly call SDMs we assessed the proportion of incorrectly phased alleles in 26 randomly chosen individuals (one from each sub-populations) using GATK’s ReadBackedPhasing tool, which uses the co-occurrence of alleles in the same read to infer phase at nearby variants. On average only 0.34% of SDMs were phased discordantly to the original data using this approach (a further 1.7% could not be phased using the read backed phasing approach alone). This number was found to be relatively consistent across population groups (0.28% in Europeans to 0.4% in Africans).

Illumina HiSeq X paired-end sequencing data for 1370 Lothian Birth Cohort (Deary et al., 2012; Taylor et al., 2018) individuals (mean sequencing depth of 36X) were aligned to the build 38 version of the human reference genome using BWA (Li and Durbin, 2009). Variants were called using GATK (DePristo et al., 2011) according to its recommended best practices. This included the use of GATKs HaplotypeCaller software that implements read based phasing of nearby alleles. After checking identities with previous array data, excluding duplicate individuals and those displaying excessive levels of heterozygosity 1358 individuals remained. All SNP coordinates were then lifted over to build 37 so as to match the 1000 genomes dataset using Crossmap (Zhao et al., 2014). Ancestral sequences available at ftp://ftp.1000genomes.ebi.ac.uk/vol1/ftp/phase1/analysis_results/supporting/ancestral_alignments/ were used to determine the ancestral alleles of each SNP, and as with the 1000 genomes data only those variants with high-confidence calls were kept (see Paten et al. (2008) for more details). SDMs were subsequently called using the same approach as the 1000 genomes data.

All SNPs were annotated using variant effect predictor (VEP, McLaren et al. (2016)) with gene models from the version 85 release of Ensembl. SDMs were subsequently reannotated using custom python scripts to correctly annotate the consequence of the double base change in coding regions.

All non-coding SNPs, MNPs and SDMs were defined with respect to their triplet context on the reference strand. This meant that all non-coding SDMs were recorded twice, once with respect to their immediate 5’ neighbour and once with respect to their 3’ neighbour (Fig 1). Non-coding SNPs were recorded with respect to all three frames within which they fell. On the other hand coding polymorphisms were recorded with respect to just the actual codon within which they occurred. This enabled the direct comparison of the occurrence of SNPs and SDMs inside annotated codons to matching triplet bases outside of coding regions.

### Calculating observed and expected numbers of SDMs

The relative occurrence of particular mutation types were calculated as in previous studies of SNPs (Harris and Pritchard, 2017). First, the number of distinct changes in a particular triplet context (m) was counted in each population (P). Then this count, C_p_(m) was converted to a mutational fraction by dividing it by the count of all observed changes. For SDMs, an individual was a carrier if at least one of their haplotypes carried both derived alleles at the corresponding pair of nucleotides.

In various analyses the two changes that comprise an SDM were separated into those that came first (i.e. the derived allele at just one of the two nucleotides is observed by itself in the population, see Fig 1) and those that came second. Their relative occurrence were also calculated as above.

The expected SDM frequencies of occurrence were derived from background SNP numbers by calculating the conditional probability of observing an SDM comprising the corresponding two changes (Eq 1).

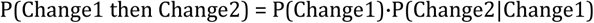
(1)
Where P(Change1) is the proportion of SNPs displaying the corresponding change when defined by their triplet context. P(Change2|Change1) is the proportion of SNPs displaying the same change as change 2 among all SNPs with the same ancestral triplet and where the change is at a position in the triplet neighbouring the location of the first change. For example, if the first change was AAA>ATA and the second change ATA>ATC, P(Change2|Change1) would correspond to the number of ATA>ATC intergenic SNPs divided by the sum of all ATA>ATB and ATA>BTA changes (where B can be any nucleotide except A). This therefore accounts for the fact that the second change in an SDM had to occur at a base neighbouring, but not at, the location of the first, and involve the triplet the first change had created.

To calculate the expected number of coding SDMs given their frequency of occurrence in intergenic regions we first corrected the counts of each intergenic SDM for the difference in triplet occurrence between coding and intergenic regions. This was done for each SDM by dividing their observed number in intergenic regions by the ratio of intergenic to coding triplet counts for the corresponding ancestral triplet. To account for the general underrepresentation of SDMs in coding regions we then divided this value for each SDM by the sum across all SDMs to get the relative, normalised mutational fractions of each change. This was finally multiplied by the total number of all coding SDMs so that the observed and expected counts were on the same scale.

### Statistical analyses

To test whether the impact of CpG dynamics on SDM numbers depended on the form of the original base change, we fit the negative binomial generalized linear model specified in Eq (2).

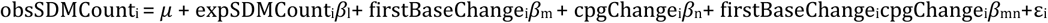
(2)
Where obsSDMCount_i_ corresponds to the observed number of SDMs of type i (e.g. AAA>ATA>ATT), expSDMCount_i_ is the expected number given background SNP mutational fractions (see Eq 1) and firstBaseChange_i_ and cpgChange_i_ are the first base change and impact of both changes on any CpG sites respectively. The significance of the interaction term was assessed using ANOVA.

Significance testing in Fig 4 and Supplementary Figure S12 was carried out as in Harris and Pritchard (2017) i.e. we used their iterative approach of undertaking conditionally independent chi-square tests to try and minimise false positive significant results. To test for the enrichment of specific codon changes among coding SDMs, having accounted for background rates of coding SNP and intergenic SDM changes, we used multiple linear regression as specified in Eq 3 to Eq 6.

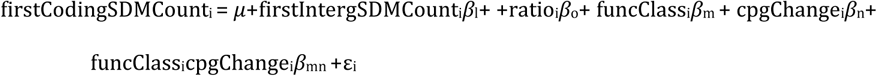
(3)

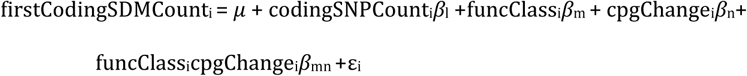
(4)

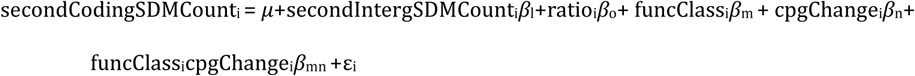
(5)

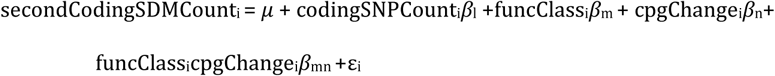
(6)
Where firstCodingSDMCount_i_ corresponds to the number of distinct first changes among coding SDMs that match change i, where each i is one of the 576 possible single base difference between two codons. secondCodingSDMCount_i_ is the corresponding count among the second changes of coding SDMs and firstIntergSDMCount_i_ and secondIntergSDMCount_i_ are the corresponding counts among the same nucleotide triplets at intergenic SDMs. codingSNPCount_i_ is the count of the same change observed among coding SNPs, funcClass_i_ is the functional impact of the corresponding change (missense, stop gained etc) and cpgChange_i_ is the impact on any CpG sites (lost or created). Ratio_i_ is the ratio of the counts of the corresponding ancestral triplet in intergenic and coding regions. Any interaction effect between the functional impact of the given codon change and its impact on CpG sites is represented by *β*_mn_. The results of these four models are shown in Supplementary Figure S16 and Supplementary Figure S17. Multiple linear regression was also used in the test for epistatic selection among coding SDMs as specified in Eq 7.

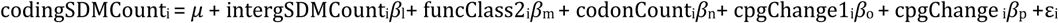
(7)
Where codingSDMCount_i_ is the number of each SDM, i, in the genome where i is restricted to the 663 SDM where the first change is missense. intergSDMCount_i_ is the count of the same SDM, i, in intergenic regions. funcClass2_i_ is the functional impact of the second change in SDM i, and cpgChange1_i_ and cpgChange2_i_ are the impact on CpG sites of the first and second changes in the SDM respectively. codonCount_i_ is the number of codons that encode the final amino acid created by the SDM, to account for the fact that amino acids with only one codon are generally favoured by SDMs. A goodness-of-fit test confirmed that the Poisson model suitably fit the data (Chi-squared p=0.997).

## Supporting information

## Acknowledgements

JGDP is supported by the Biotechnology and Biological Sciences Research Council (BBSRC, Grant No. BBS/E/D/10002071) and the whole genome sequencing of the Lothian Birth Cohorts was funded through an institutional award to the Roslin Institute from the BBSRC. The Lothian Birth Cohorts are supported by Age UK (Disconnected Mind programme). IJD is supported by the Centre for Cognitive Ageing and Cognitive Epidemiology, which is funded by the Medical Research Council and the Biotechnology and Biological Sciences Research Council (Grant No. MR/K026992/1). We thank the Lothian Birth Cohorts’ participants and research team for their help.

## Supporting information

**Supplementary Figure S1** Number of SNPs and SDMs each individual carries

**Supplementary Figure S2** African population SDM (red) and SNP (blue) densities in 1Mb windows across the genome **Supplementary Figure S3** American population SDM (red) and SNP (blue) densities in 1Mb windows across the genome **Supplementary Figure S4** East Asian population SDM (red) and SNP (blue) densities in 1Mb windows across the genome **Supplementary Figure S5** European population SDM (red) and SNP (blue) densities in 1Mb windows across the genome **Supplementary Figure S6** South Asian population SDM (red) and SNP (blue) densities in 1Mb windows across the genome **Supplementary Figure S7** SDM versus SNP densities in 1Mb windows along the genome. Densities in different population groups are shown and the outlier windows corresponding to the MHC region on chromosome 6 are indicated.

**Supplementary Figure S8** Counts of SDMs by change (rows) and haplotype frequencies (columns). Haplotype frequencies are broken down into three heatmaps by number of ancestral alleles, and the bar chart along the top of each heatmap indicates the total number of SDMs with the corresponding haplotype frequency.

**Supplementary Figure S9** The same as Figure 2D but without restricting to sites where the ancestral haplotype could be determined and folding together changes involving the same three haplotypes.

**Supplementary Figure S10** The same as Figure 2D but for MNPs rather than SDMs.

**Supplementary Figure S11** Principal component analysis of individuals according to the difference in the mutational fraction of first and second changes among SDMs. The mutational fraction of each second change in an SDM defined by their base change and triplet nucleotide context was subtracted from the corresponding mutational fraction of first changes. The resulting differences were then used as input to the PCA. The continental groups still separate in this analysis suggesting the first and second changes display different biases between populations.

**Supplementary Figure S12** Same as Fig 4 but for the Lothian Birth Cohort genomes.

**Supplementary Figure S13** Frequencies and counts of a subset of intergenic SDMs and SNPs across the 1000 genomes cohort. The three leftmost heatmaps indicate allele frequencies of the intergenic SDMs where the second change is CpG → TpG, i.e. the potential deamination of a CpG site. The numbers of intergenic SDMs of the corresponding allele frequency are indicated by the intensity of colour. The derived allele frequencies and numbers of intergenic SNPs matching each constitutive change are indicated in the final two heatmaps. The original base change in the SDM and its impact on any CpG sites is also indicated (the base change being orientated with respect to the CpG → TpG change). The yellow histograms represent the total counts of each corresponding intergenic SDM or SNP.

**Supplementary Figure S14** The occurrence of all SDMs in a 6mer context involving the gain and loss of CpG sites in the 1000 genomes cohort normalised by the frequency of the ancestral 6mer in the genome. SDMs involving the same nucleotide changes are grouped by colour. SDMs involving a CA>CG>TG change or its reverse complement are common in largely all 6mer contexts.

**Supplementary Figure S15** The importance of CpG dynamics in shaping SDM numbers depends on the form of the original base change. Each dot represents the observed count of a particular form of SDM. Filled circles and error bars represent the expected number of SDM of the given category having controlled for background rates of change at SDM. These expected numbers were generated from prediction outputs of Eq 2 when the expected number of SDM variable was fixed at its median observed value. The interaction term in Eq 2, assessed using ANOVA, was significant (p < 2.2×10^−16^) in both cohorts. **Supplementary Figure S16** The enrichment and depletion of changes in coding SDMs having accounted for the corresponding occurrence of changes observed among SNPs. The observed number of each codon change among the first and second change of coding SDMs was fitted against its consequence, effect on CpG sites and observed occurrence among SNPs. See Eq 4 and Eq 6 in the methods section for more details.

**Supplementary Figure S17** The enrichment and depletion of changes in coding SDMs having accounted for the corresponding frequency of occurrence of changes observed among intergenic SDMs. The observed number of each codon change among the first and second change of coding SDMs was fitted against its consequence, effect on CpG sites and observed frequency of occurrence among intergenic SDMs. See Eq 3 and Eq 5 in the methods section for more details.

**Supplementary Figure S18** Testing for epistatic selection across neighbouring, coding nucleotide changes. Restricting the analysis to SDMs where the first change leads to a missense change in coding regions we tested whether the frequency of occurrence of these SDMs was related to the consequence of the second change. We controlled for the number of codons that encode the final amino acid, the impact of the second change in the SDM on CpG sites and the corresponding frequency of occurrence of the first change at coding SNPs in this analysis. See Eq 7 in the methods section for more details.

