## Supplementary material for "Linked mutations at adjacent nucleotides have shaped human population differentiation and protein evolution"

**Supplementary Figure S1** Number of SNPs and SDMs each individual in the 1000 genomes cohort carries.

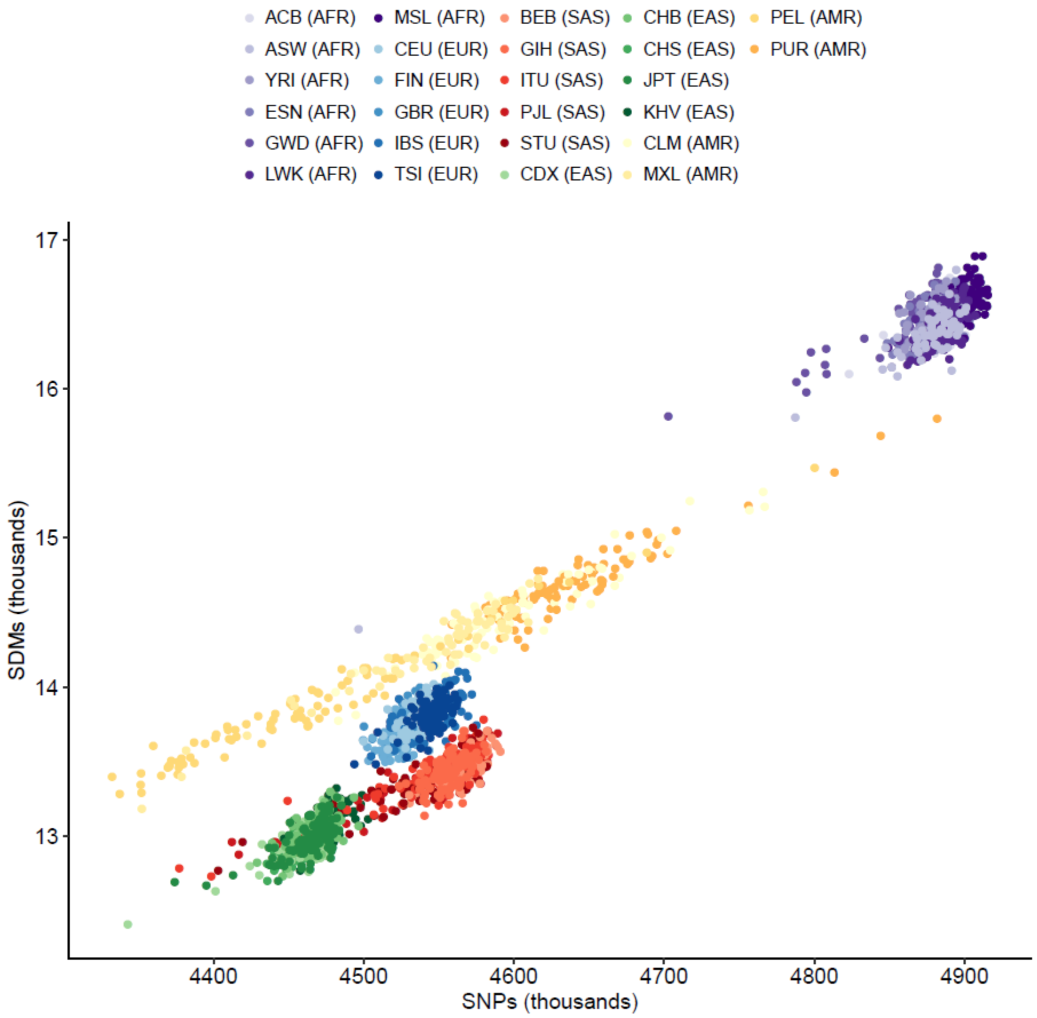

**Supplementary Figure S2** African population SDM (red) and SNP (blue) densities in 1Mb windows across the genome

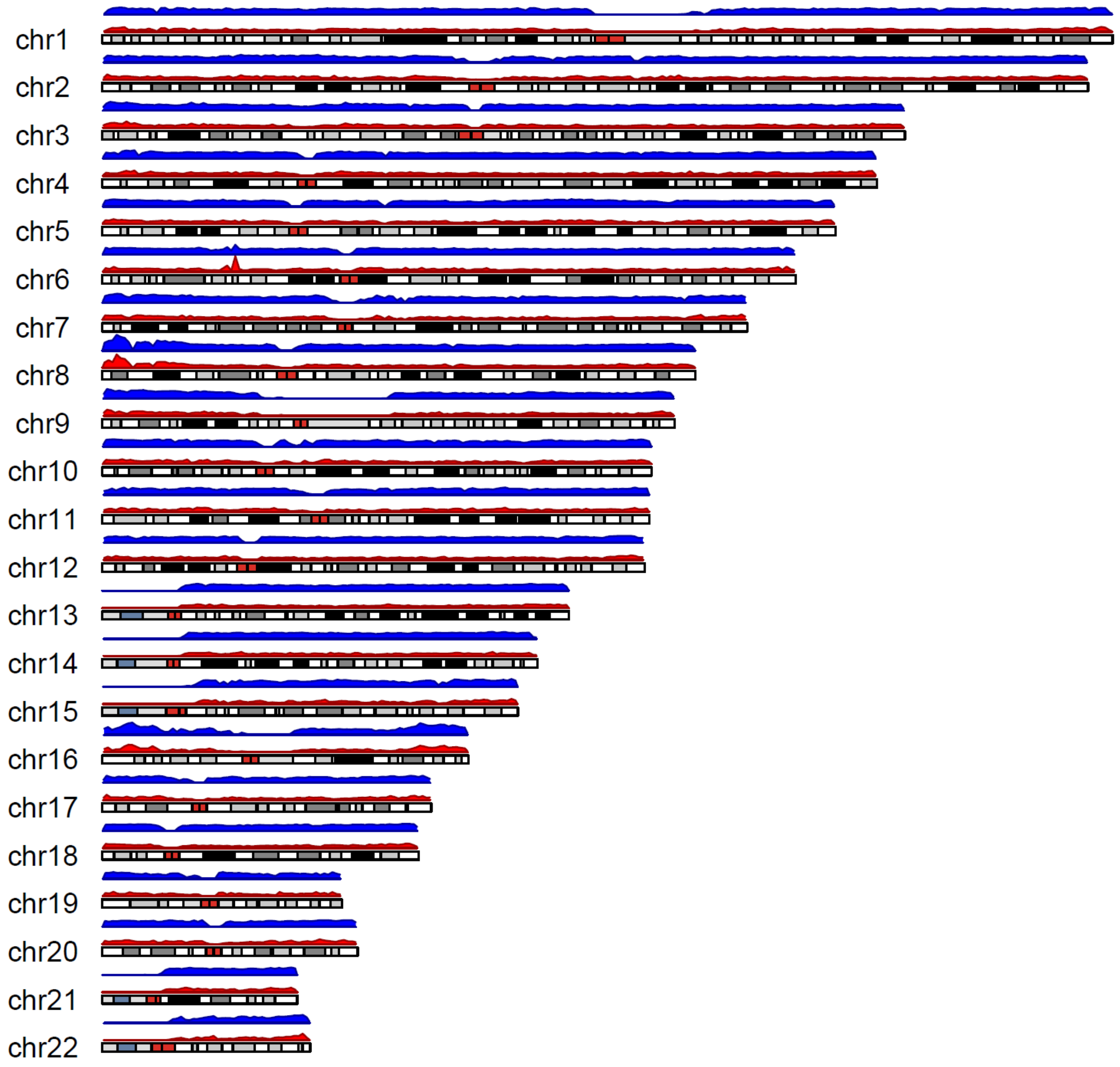

**Supplementary Figure S3** American population SDM (red) and SNP (blue) densities in 1Mb windows across the genome

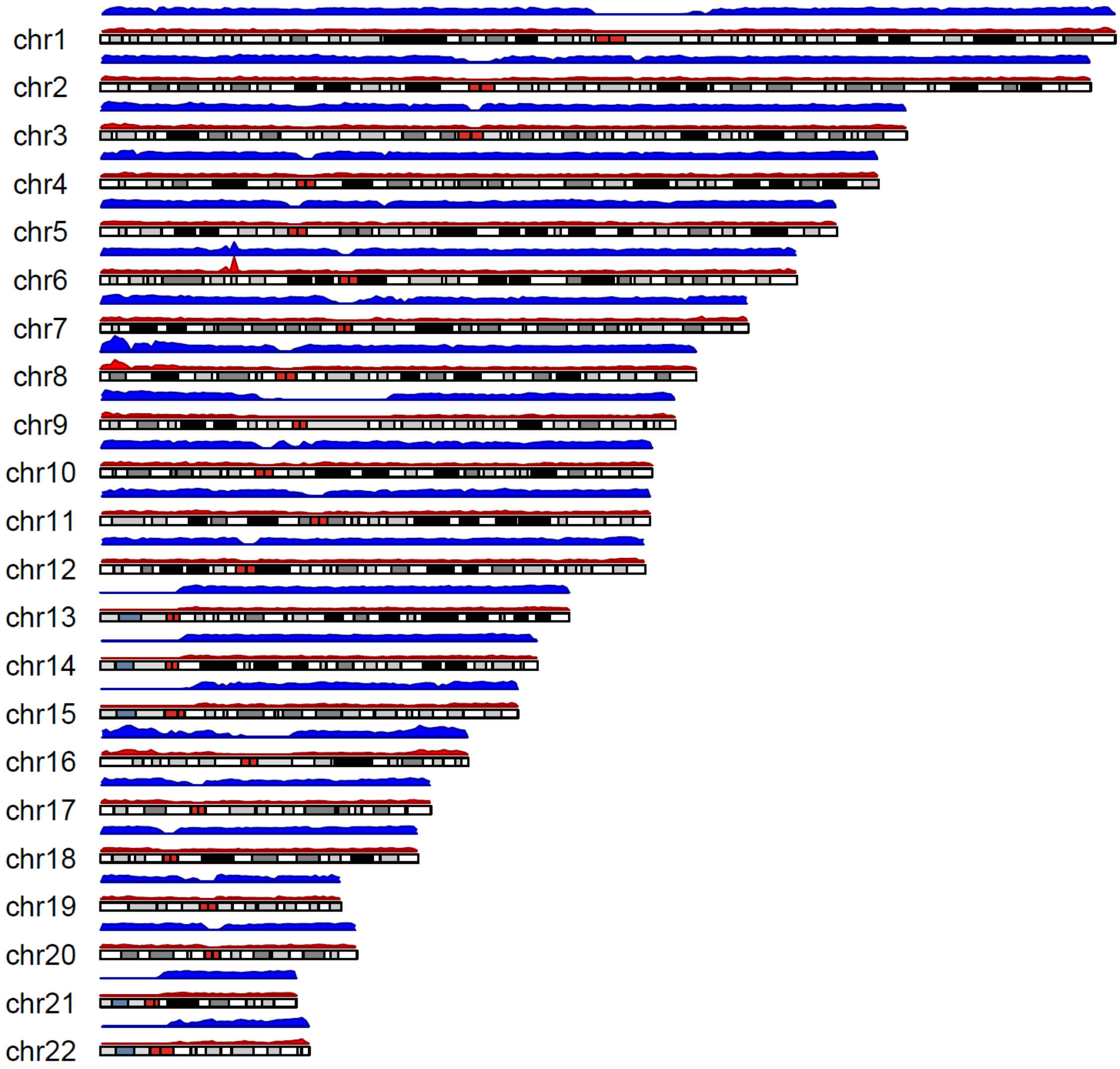

**Supplementary Figure S4** East Asian population SDM (red) and SNP (blue) densities in 1Mb windows across the genome

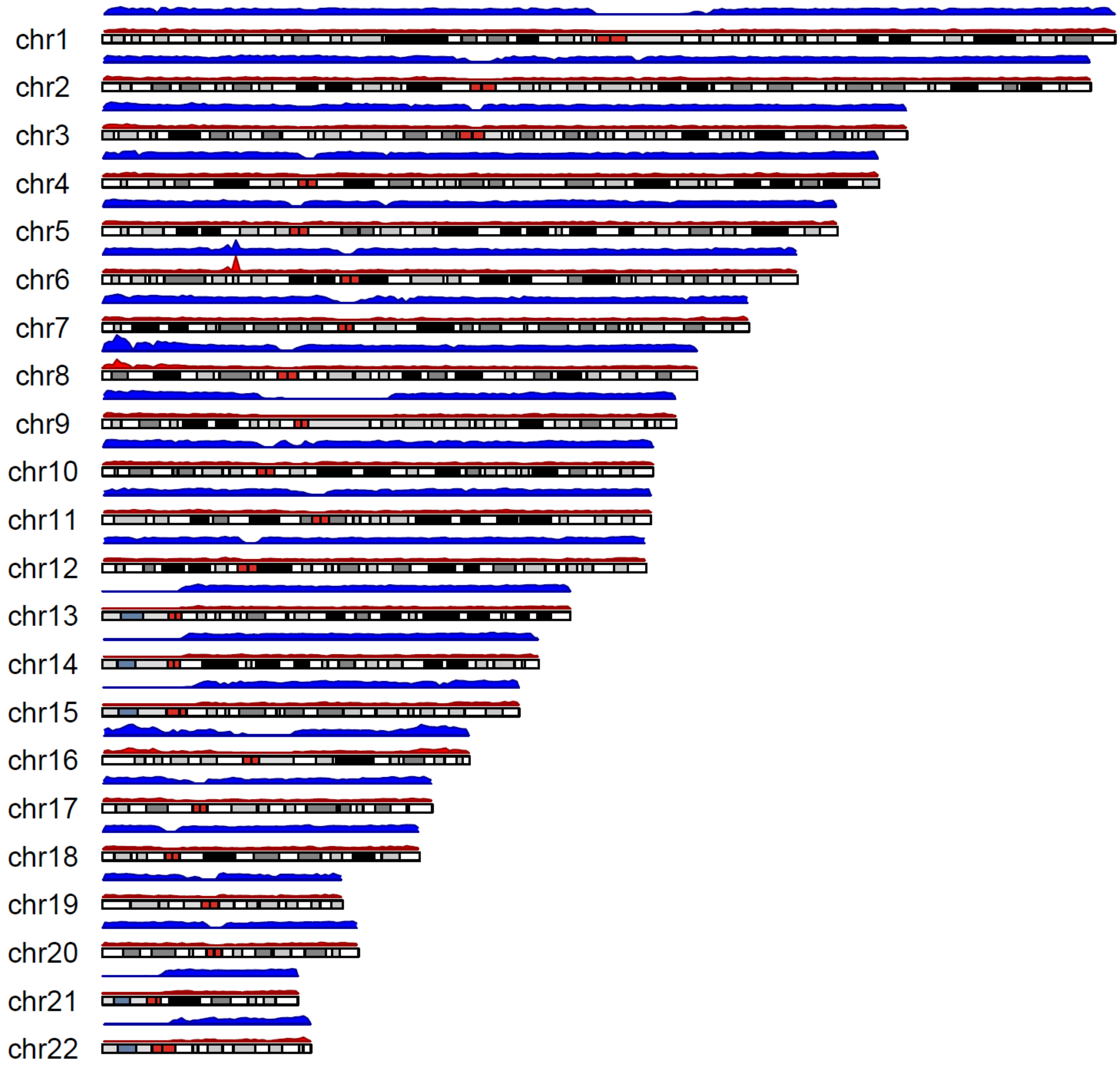

**Supplementary Figure S5** European population SDM (red) and SNP (blue) densities in 1Mb windows across the genome

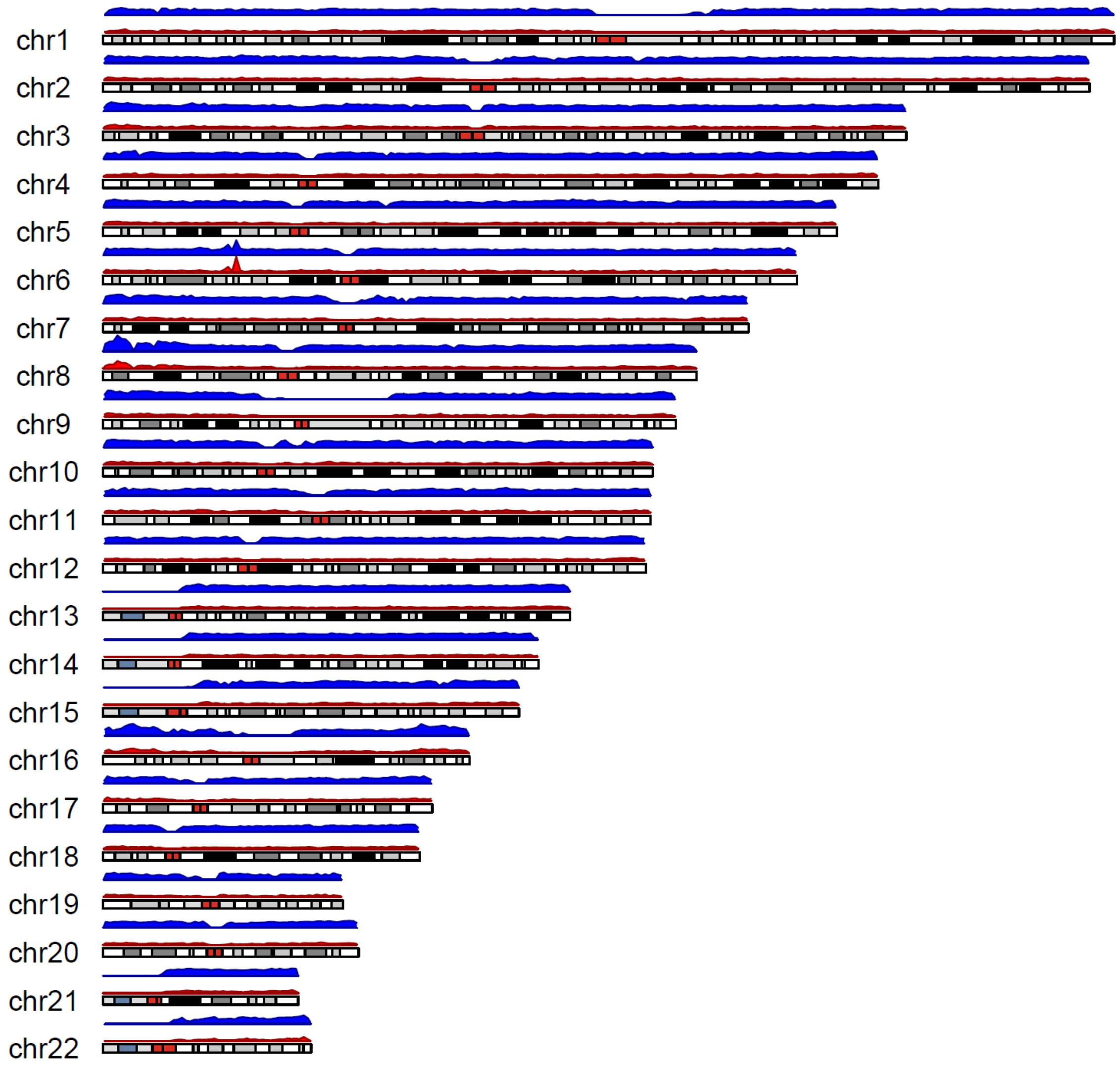

**Supplementary Figure S6** South Asian population SDM (red) and SNP (blue) densities in 1Mb windows across the genome

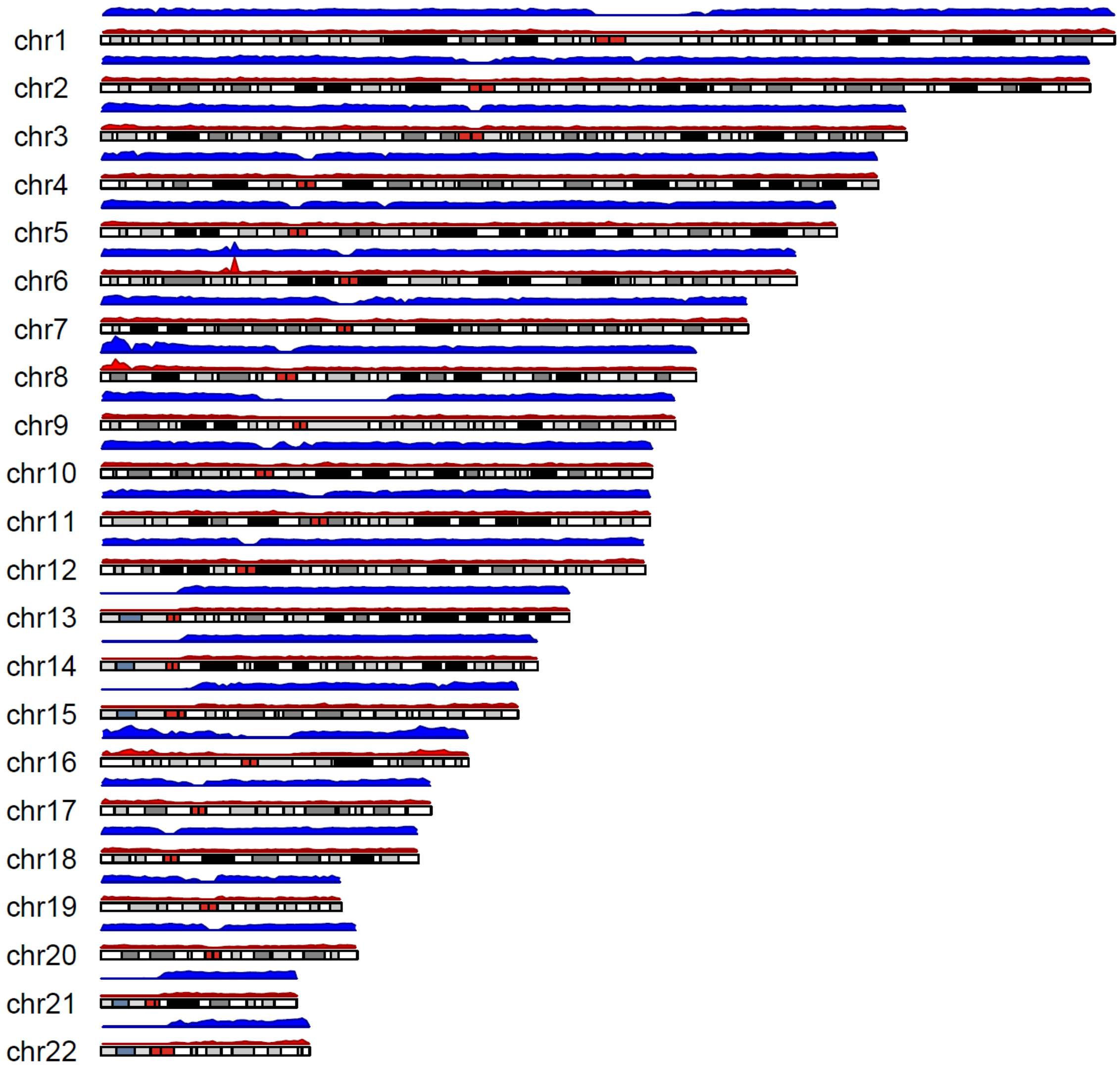

**Supplementary Figure S7** SDM versus SNP densities in 1Mb windows along the genome. Densities in different population groups are shown and the outlier windows corresponding to the MHC region on chromosome 6 are indicated.

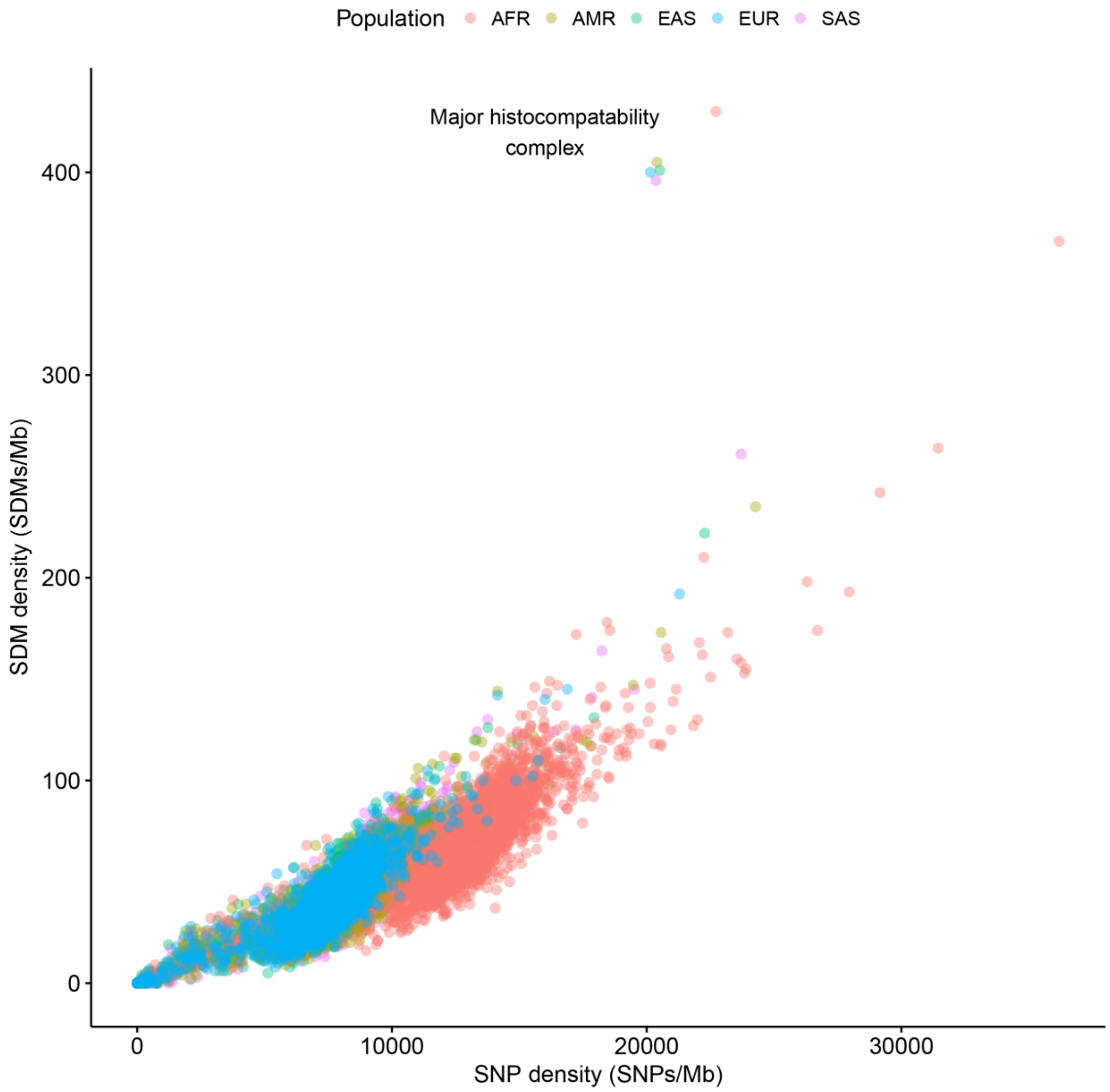

**Supplementary Figure S9** The same as Figure 2D but without restricting to sites where the ancestral haplotype could be determined and folding together changes involving the same three haplotypes.

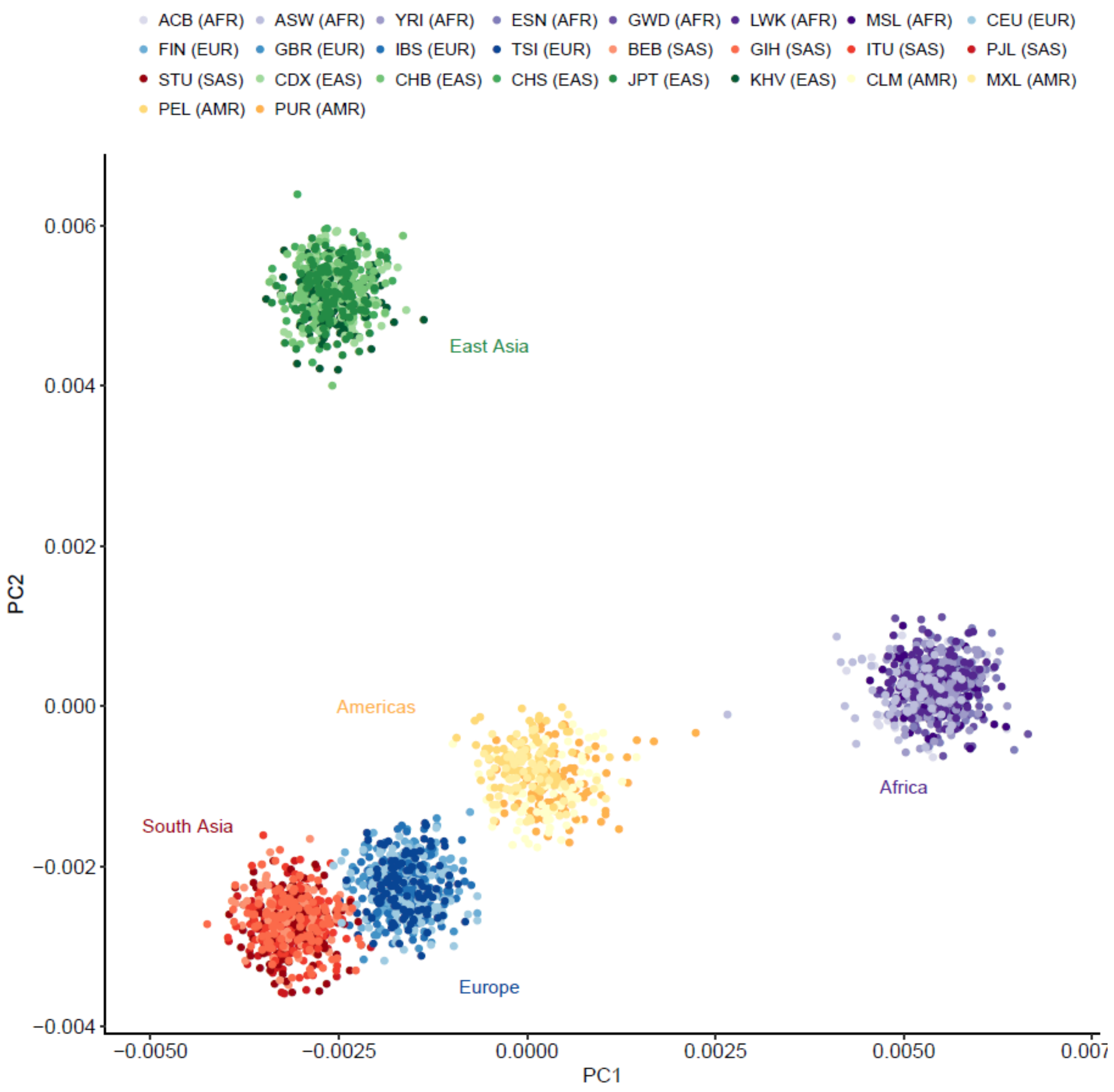

**Supplementary Figure S10** The same as Figure 2D but for MNPs rather than SDMs.

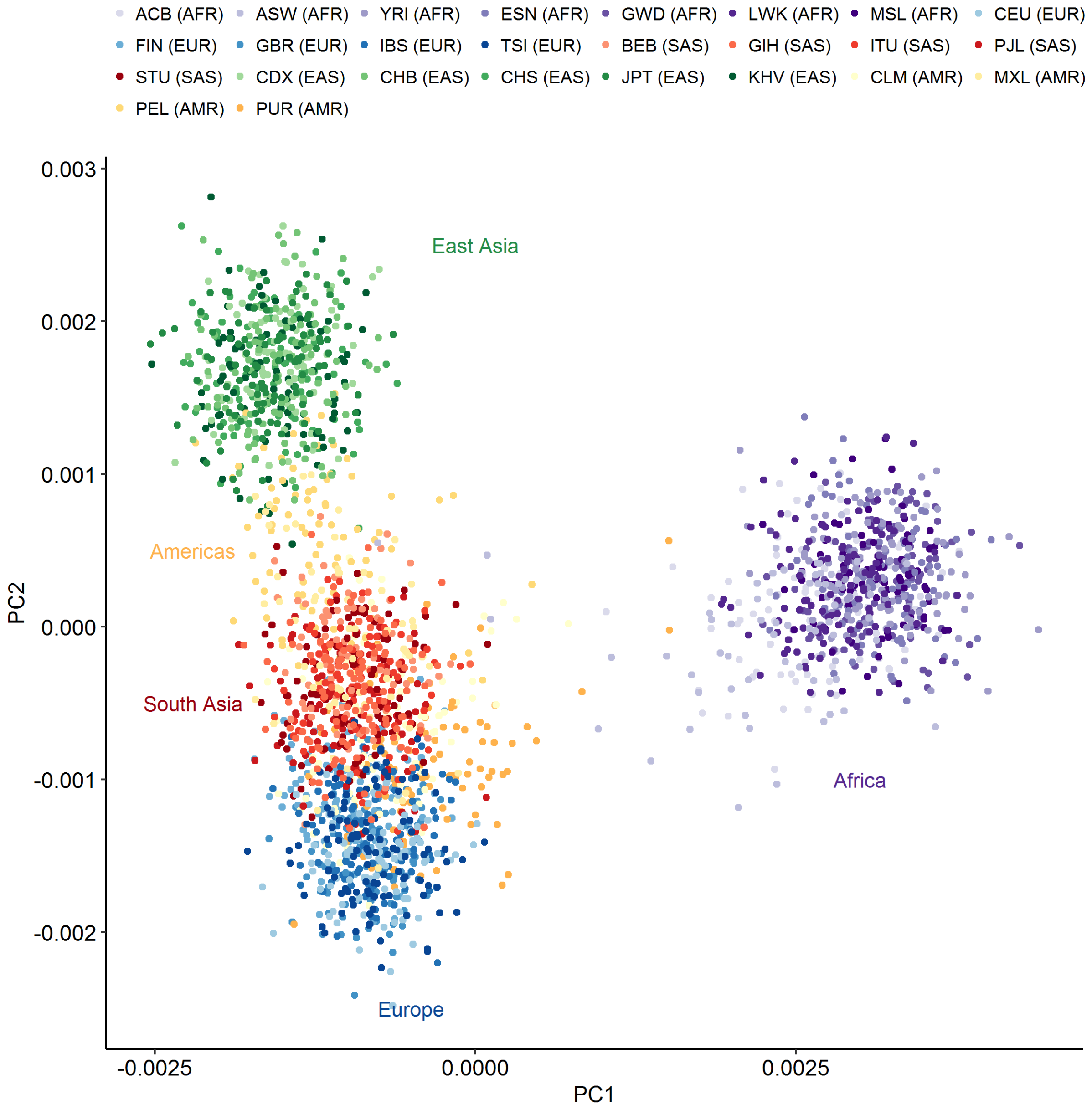

**Supplementary Figure S11** Principal component analysis of individuals according to the difference in the mutational fraction of first and second changes among SDMs. The mutational fraction of each second change in an SDM defined by their base change and triplet nucleotide context was subtracted from the corresponding mutational fraction of first changes. The resulting differences were then used as input to the PCA. The continental groups still separate in this analysis suggesting the first and second changes display different biases between populations.

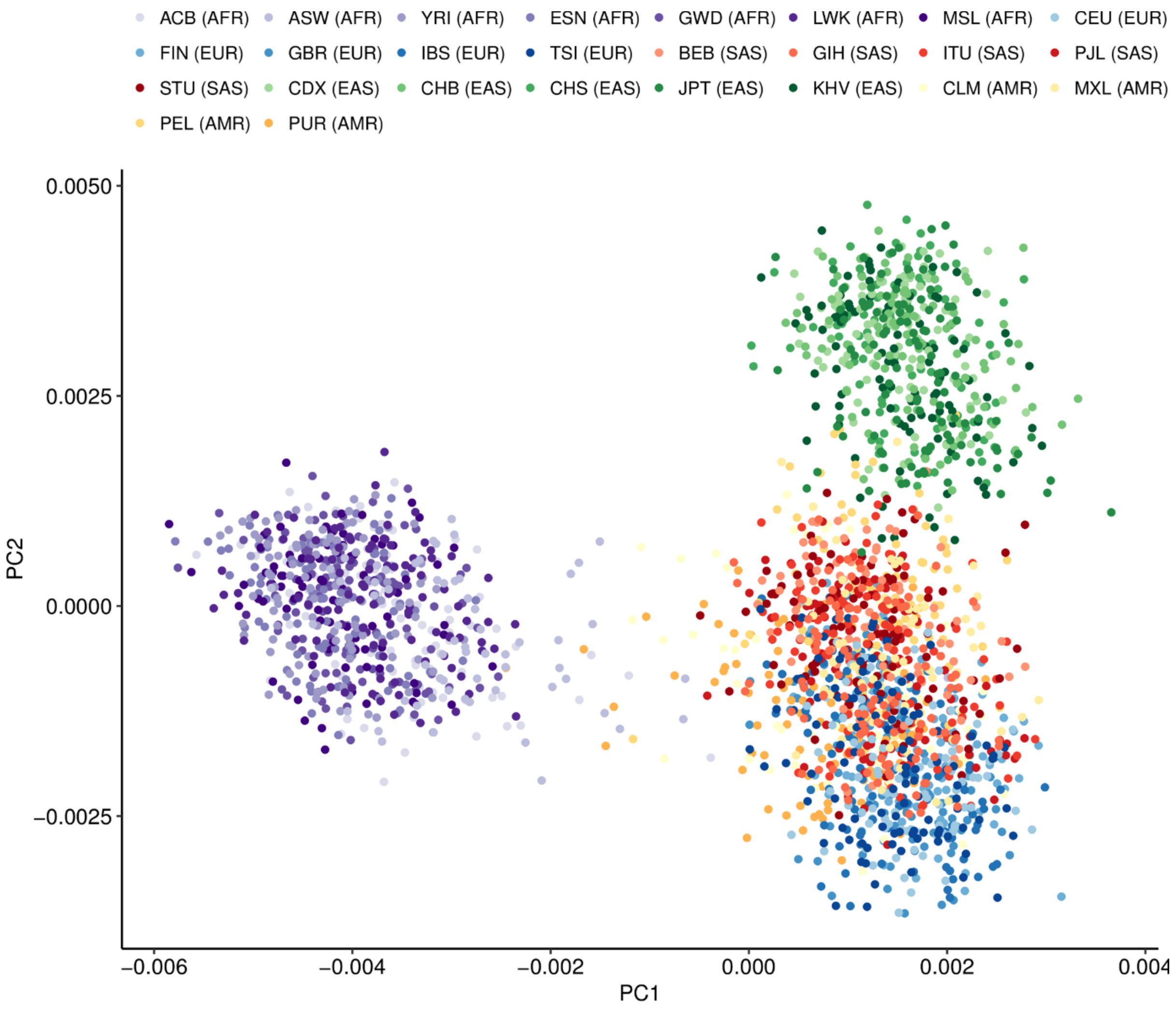

**Supplementary Figure S12** Same as Fig 4 but for the Lothian Birth Cohort genomes.

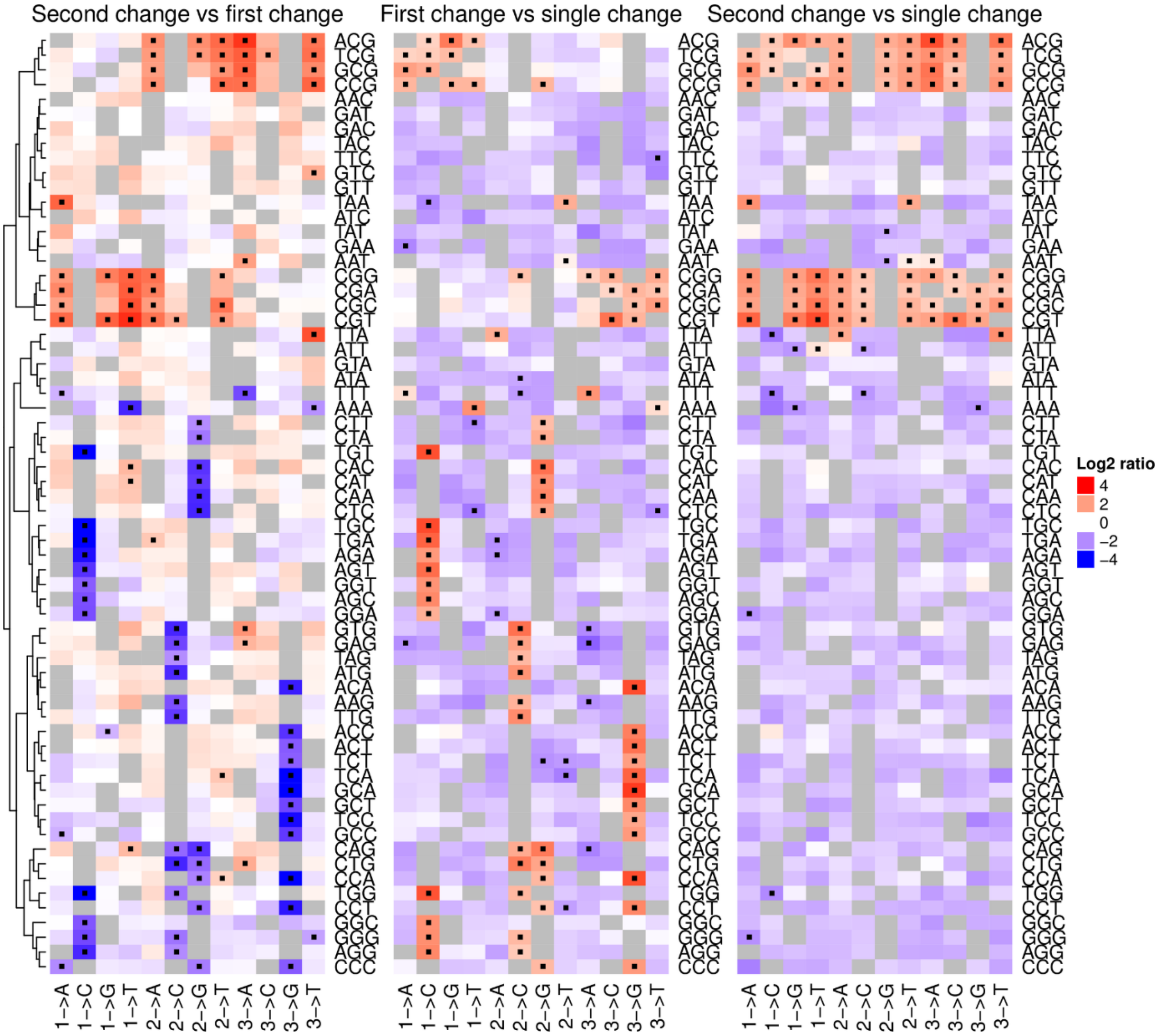

**Supplementary Figure S13** Frequencies and counts of a subset of intergenic SDMs and SNPs across the 1000 genomes cohort. The three leftmost heatmaps indicate allele frequencies of the intergenic SDMs where the second change is CpG → TpG, i.e. the potential deamination of a CpG site. The numbers of intergenic SDMs of the corresponding allele frequency are indicated by the intensity of colour. The derived allele frequencies and numbers of intergenic SNPs matching each constitutive change are indicated in the final two heatmaps. The original base change in the SDM and its impact on any CpG sites is also indicated (the base change being orientated with respect to the CpG → TpG change). The yellow histograms represent the total counts of each corresponding intergenic SDM or SNP.

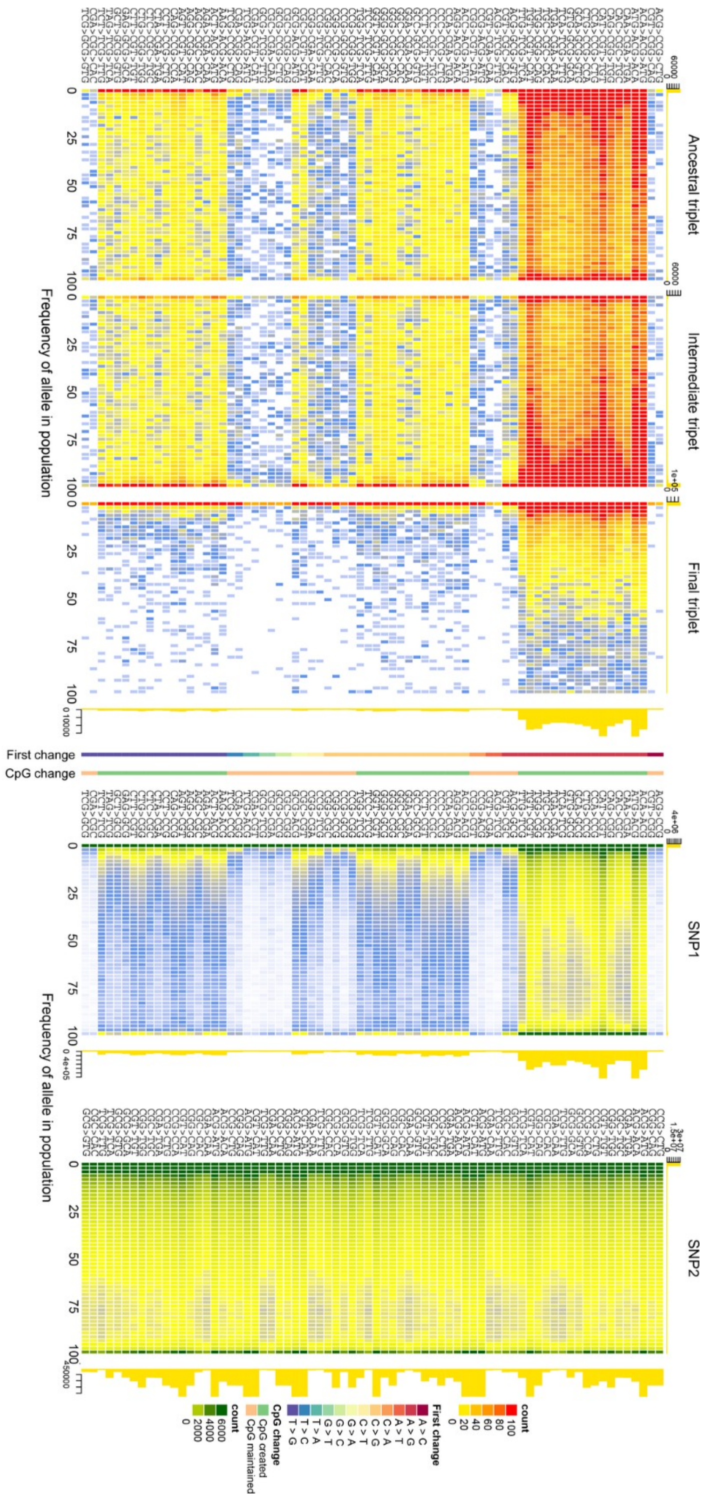

**Supplementary Figure S14** The occurrence of all SDMs in a 6mer context involving the gain and loss of CpG sites in the 1000 genomes cohort normalised by the frequency of the ancestral 6mer in the genome. SDMs involving the same nucleotide changes are grouped by colour. SDMs involving a CA>CG>TG change or its reverse complement are common in largely all 6mer contexts.

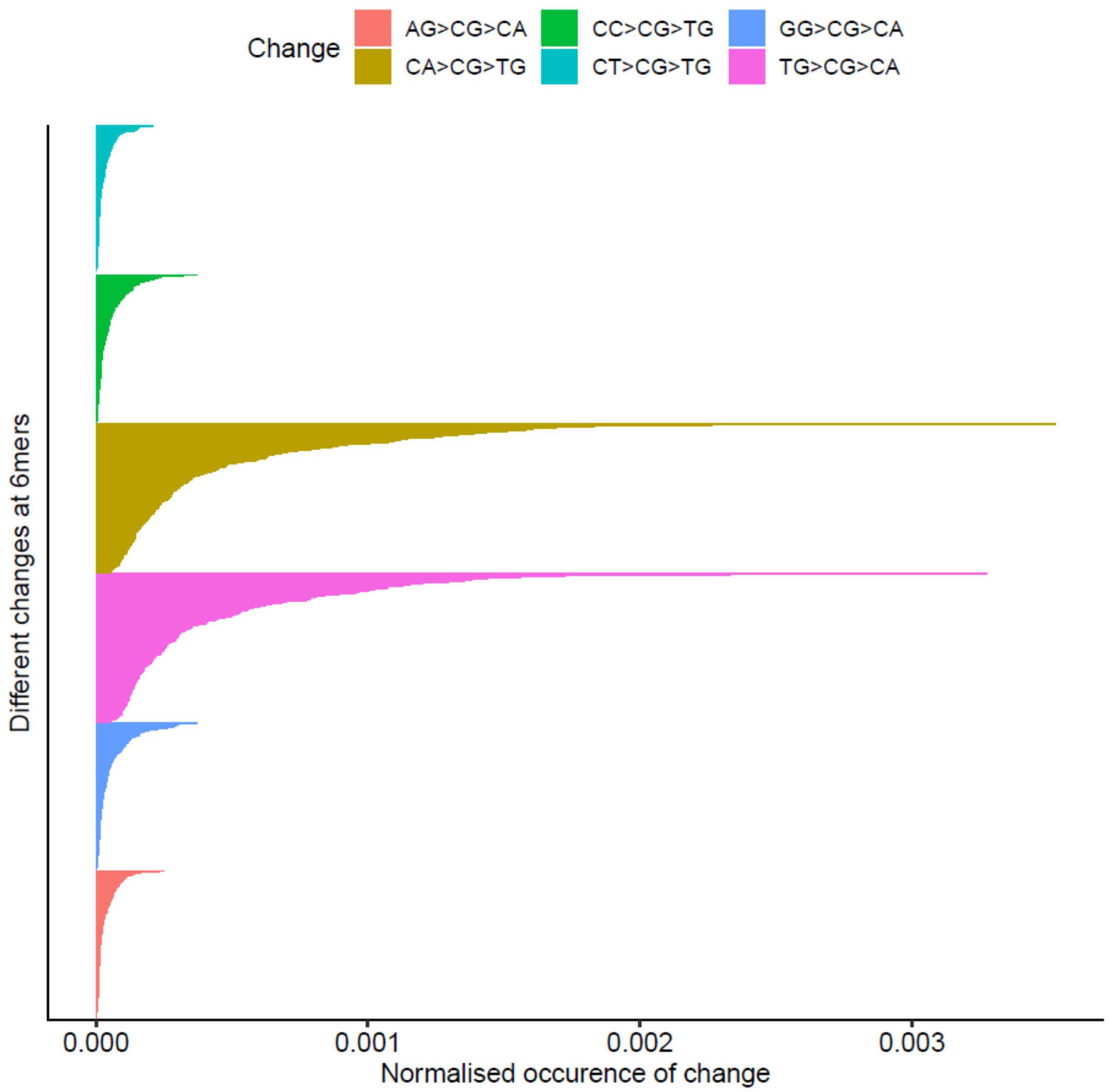

**Supplementary Figure S15** The importance of CpG dynamics in shaping SDM numbers depends on the form of the original base change. Each dot represents the observed count of a particular form of SDM. Filled circles and error bars represent the expected number of SDM of the given category having controlled for background rates of change at SDM. These expected numbers were generated from prediction outputs of Eq 2 when the expected number of SDM variable was fixed at its median observed value. The interaction term in Eq 2, assessed using ANOVA, was significant ($p < 2.2\text{x}10^{-16}$) in both cohorts.

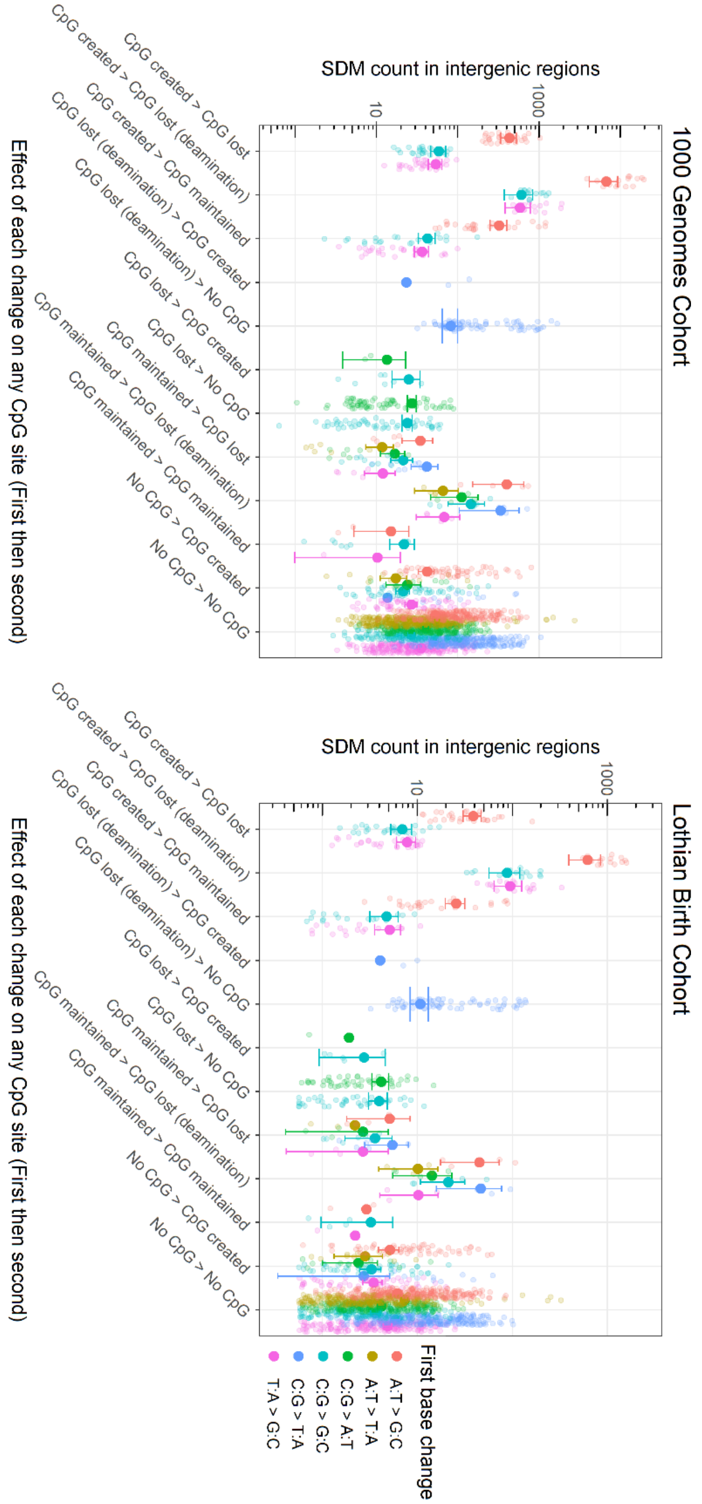

**Supplementary Figure S16** The enrichment and depletion of changes in coding SDMs having accounted for the corresponding occurrence of changes observed among SNPs. The observed number of each codon change among the first and second change of coding SDMs was fitted against its consequence, effect on CpG sites and observed occurrence among SNPs. See Eq 4 and Eq 6 in the methods section for more details.

| | *Dependent variable:* | |
|---|---|---|
| | first variant in coding MNP | second variant in coding MNP |
| single coding variant in SNP | **0.001*** ** | **0.0004*** ** |
| | **(0.0001)** | **(0.0001)** |
| **Consequence (reference level: synonymous)** | | |
| missense | 0.79 | 1.01** |
| | (0.58) | (0.43) |
| Met/start gain | 0.33 | 0.12 |
| | (1.76) | (1.30) |
| Met/start lost | 0.26 | 0.93 |
| | (1.76) | (1.30) |
| stop gain | 0.96 | 0.76 |
| | (1.18) | (0.87) |
| stop lost | 1.24 | 0.37 |
| | (1.19) | (0.88) |
| **CpG change (reference level: no CpG)** | | |
| lost | 1.10 | 1.05 |
| | (1.60) | (1.18) |
| lost (deamination) | 6.27** | **18.59*** ** |
| | (2.60) | **(1.92)** |
| made | **10.69*** ** | 0.03 |
| | **(1.37)** | (1.01) |
| maintained | 3.79** | 0.87 |
| | (1.47) | (1.09) |
| **Interactions (consequence and CpG change)** | | |
| missense:lost | -0.58 | -1.37 |
| | (1.95) | (1.44) |
| stop gain:lost | -0.93 | -1.48 |
| | (5.15) | (3.80) |
| missense:lost (deamination) | -5.18* | **12.76*** ** |
| | (2.88) | **(2.13)** |
| Met/start gain:lost (deamination) | -0.61 | **66.23*** ** |
| | (5.63) | **(4.15)** |
| stop gain:lost (deamination) | -7.76 | -9.61** |
| | (5.49) | (4.05) |
| missense:made | **-5.13*** ** | 0.23 |
| | **(1.66)** | (1.22) |
| Met/start lost:made | **25.28*** ** | 2.68 |
| | **(5.27)** | (3.89) |
| stop lost:made | -10.73** | -0.045 |
| | (3.79) | (2.80) |
| missense:maintained | -2.54 | -0.72 |
| | (1.97) | (1.45) |
| Constant | -1.27** | -0.39 |
| | (0.57) | (0.42) |
| Observations | 576 | 576 |
| $R^2$ | 0.36 | 0.76 |
| Adjusted $R^2$ | 0.34 | 0.75 |
| Residual Std. Error (df = 556) | 4.78 | 3.53 |
| F Statistic (df = 19; 556) | 16.42*** | 93.52*** |
| *Note:* | | *p<0.1; **p<0.05; ***p<0.0026 |

**Supplementary Figure S17** The enrichment and depletion of changes in coding SDMs having accounted for the corresponding frequency of occurrence of changes observed among intergenic SDMs. The observed number of each codon change among the first and second change of coding SDMs was fitted against its consequence, effect on CpG sites and observed frequency of occurrence among intergenic SDMs. See Eq 3 and Eq 5 in the methods section for more details.

| | *Dependent variable:* | |
|---|---|---|
| | first variant in coding SDM | second variant in coding SDM |
| first variant in intergenic SDM | **0.002*** ** | |
| | **(0.0001)** | |
| second variant in intergenic SDM | | **0.002*** ** |
| | | **(0.0002)** |
| Relative frequency of triplet in coding and non-coding regions | -0.002 | -0.002 |
| | (0.002) | (0.002) |
| **Consequence (reference level: synonymous)** | | |
| missense | -0.78 | 0.20 |
| | (0.49) | (0.39) |
| Met/start gain | -0.40 | -0.26 |
| | (1.52) | (1.23) |
| Met/start lost | -0.82 | 0.14 |
| | (1.52) | (1.23) |
| stop gain | -1.51 | -0.44 |
| | (1.00) | (0.80) |
| stop lost | 0.049 | 0.56 |
| | (1.80) | (1.45) |
| **CpG change (reference level: no CpG)** | | |
| lost | -0.44 | -0.35 |
| | (1.38) | (1.11) |
| lost (deamination) | **12.79*** ** | -6.34* |
| | **(2.11)** | (3.33) |
| made | 2.67** | -0.19 |
| | (1.25) | (0.95) |
| maintained | 2.00 | -0.25 |
| | (1.27) | (1.02) |
| **Interactions (consequence and CpG change)** | | |
| missense:lost | 0.65 | -0.77 |
| | (1.69) | (1.36) |
| stop gain:lost | 0.88 | -0.63 |
| | (4.45) | (3.58) |
| missense:lost (deamination) | -5.13** | **12.09*** ** |
| | (2.50) | **(2.01)** |
| Met/start gain:lost (deamination) | -2.34 | **58.35*** ** |
| | (4.87) | **(4.00)** |
| stop gain:lost (deamination) | -11.74** | -0.75 |
| | (4.73) | (3.96) |
| missense:made | **-5.71*** ** | 0.35 |
| | **(1.43)** | (1.15) |
| Met/start lost:made | -2.31 | 1.24 |
| | (4.90) | (3.67) |
| stop lost:made | **-11.91*** ** | 0.24 |
| | **(3.29)** | (2.64) |
| missense:maintained | -1.29 | -0.06 |
| | (1.70) | (1.37) |
| Constant | 0.94 | 0.23 |
| | (0.44) | (0.36) |
| Observations | 576 | 576 |
| $R^2$ | 0.52 | 0.79 |
| Adjusted $R^2$ | 0.50 | 0.78 |
| Residual Std. Error (df = 555) | 4.14 | 3.33 |
| F Statistic (df = 20; 555) | 30.21*** | 103.21*** |
| *Note:* | | *p<0.1; **p<0.05; ***p<0.0026 |

**Supplementary Figure S18** Testing for epistatic selection across neighbouring, coding nucleotide changes. Restricting the analysis to SDMs where the first change leads to a missense change in coding regions we tested whether the frequency of occurrence of these SDMs was related to the consequence of the second change. We controlled for the number of codons that encode the final amino acid, the impact of the second change in the MNP on CpG sites and the corresponding frequency of occurrence of the first change at coding SNPs in this analysis. See Eq 7 in the methods section for more details.

| | *Dependent variable:* |
|---|---|
| | number of SDMs in coding regions (only SDMs where first change is missense) |
| Number of coding SNPs matching first change | **0.00026*** ** |
| | **(0.0000095)** |
| **Consequence of second change (reference level: synonymous)** | |
| missense | **-0.32*** ** |
| | **(0.099)** |
| Met/start gain | **-1.41*** ** |
| | **(0.29)** |
| stop gain | -0.51 |
| | (0.29) |
| reversion | -0.14 |
| | (451) |
| **Number of codons for final amino acid (reference level: 1** | |
| 2 | **-0.88*** ** |
| | **(0.16)** |
| 3 | **-0.82*** ** |
| | **(0.27)** |
| 4 | **-0.87*** ** |
| | **(0.17)** |
| 6 | **-1.24*** ** |
| | **(0.18)** |
| **CpG impact of second change (reference level: no CpG)** | |
| lost | 0.46** |
| | (0.21) |
| lost (deamination) | **3.2*** ** |
| | **(0.098)** |
| made | -0.43* |
| | (0.25) |
| maintained | 0.26 |
| | (0.25) |
| Constant | -0.74 |
| | (0.20) |
| Observations | 1632 |
| Log Likelihood | -1127.65 |
| Akaike Inf. Crit. | 2,283.30 |
| *Note:* | *p<0.1; **p<0.05; ***p<0.0033 |
