## Supplementary figures and images for "Linked mutations at adjacent nucleotides have shaped human population differentiation and protein evolution"

### Supplementary file 3

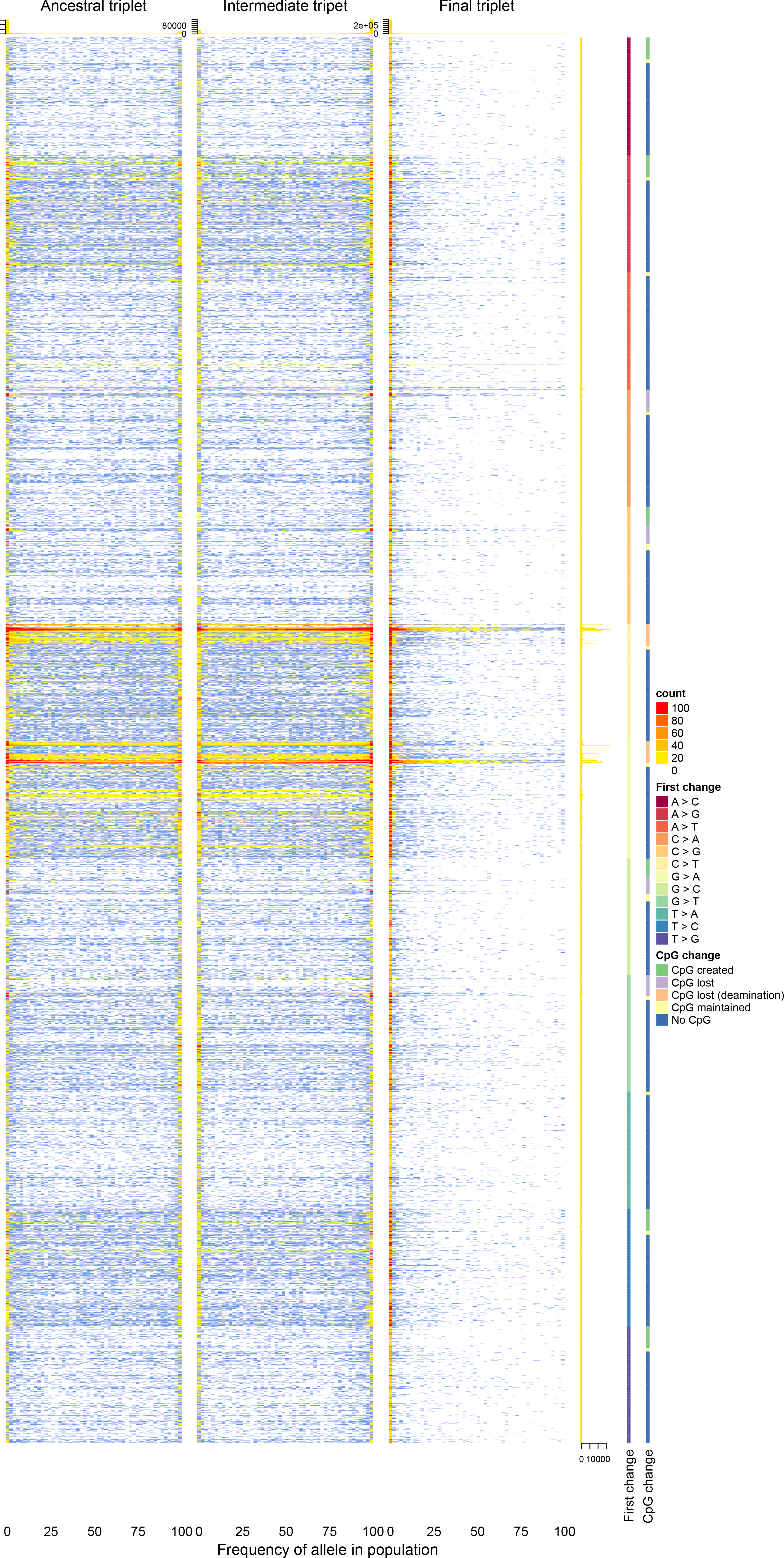
